# Actin enhances ER-mitochondrial calcium transfer and IMM constriction during mitochondrial division

**DOI:** 10.1101/192674

**Authors:** Rajarshi Chakrabarti, Wei-Ke Ji, Radu V. Stan, Jaime de Juan Sanz, Timothy A. Ryan, Henry N. Higgs

## Abstract

Mitochondrial division requires division of both the inner and outer mitochondrial membranes (IMM and OMM, respectively). Interaction with endoplasmic reticulum (ER) promotes OMM division by recruitment of the dynamin Drp1, but effects on IMM division are not well characterized. We previously showed that actin polymerization through the ER-bound formin INF2 stimulates Drp1 recruitment in mammalian cells. Here, we show that INF2-mediated actin polymerization stimulates a second mitochondrial response independent of Drp1: a rise in mitochondrial matrix calcium through the mitochondrial calcium uniporter. ER stores supply the increased mitochondrial calcium, and the role of actin is to increase ER-mitochondria contact. Myosin IIA is also required for this mitochondrial calcium increase. Elevated mitochondrial calcium in turn activates IMM constriction in a Drp1-independent manner. IMM constriction requires electron transport chain activity. IMM division precedes OMM division. These results demonstrate that actin polymerization independently stimulates the dynamics of both membranes during mitochondrial division: IMM through increased matrix calcium, and OMM through Drp1 recruitment.

## Introduction

Contacts between the endoplasmic reticulum (ER) and mitochondrion are now widely appreciated as important for communication in several respects, including lipid synthesis and calcium transfer (Phillips and Voeltz, 2016). For calcium transfer, close ER-mitochondrial contacts (<30 nm) enable a disproportionate amount of stimulus-induced calcium release from ER to be taken up by mitochondria instead of being released into the cytosol (Rizzuto et al., 1998; Csordás at al., 2006; Giacomello et al., 2010; Csordás at al., 2010). This up-take is mediated by the mitochondrial calcium uniporter, MCU (Baughman et al., 2011; De Stefani et al., 2011). Moderate increases in mitochondrial calcium stimulate oxidative catabolism through activation of several dehydrogenases (Denton et al., 2009), whereas excessive mitochondrial calcium can trigger apoptosis (Baffy et al., 1993). While a number of proteins have been shown to mediate ER-mitochondrial contacts (Phillips and Voeltz, 2016), mechanisms controlling these contacts during cell stimulation are unclear.

ER-mitochondrial contact also stimulates mitochondrial division (Friedman et al., 2011), which is required for diverse aspects of normal cellular physiology, including proper distribution of mitochondrial genomes (Lewis et al., 2016), metabolic adaptation (Mishra and Chan, 2016), mitophagy (Youle and van der Bliek, 2012; Burman et al., 2017) and immune response (Pernas and Scorano, 2016). Defects in mitochondrial division link to multiple pathologies, particularly neurodegenerative diseases (Nunnari and Suomalainen, 2012). Most mechanistic focus has been on outer mitochondrial membrane (OMM) division, with the dynamin GTPase Drp1 being a key factor (Labrousse et al, 1999; Labbe et al., 2014). Drp1 oligomerizes into a ring encircling the OMM, and GTP hydrolysis by Drp1 drives OMM constriction, leading to division. We have shown that one effector of ER-stimulated mitochondrial division in mammals is the ER-bound formin INF2, with INF2-mediated actin polymerization playing a key role in Drp1 recruitment to and oligomerization at division sites (Korobova et al., 2013; Ji et al., 2015). INF2 is linked to two human diseases: the neuropathy Charcot-Marie-Tooth disease (Boyer at al., 2011) and the kidney disease focal segmental glomerulosclerosis (Brown et al., 2010). Myosin II is also required for this process (DuBoff et al., 2012; Korobova et al., 2014), as well as the mitochondrially-bound actin polymerization factor Spire1C (Manor et al., 2015). Other mechanisms of actin polymerization might also stimulate mitochondrial division (Li et al., 2015; Moore et al., 2016). Recent work shows that dynamin 2 accumulates on the OMM subsequent to Drp1, and acts at later stages of mitochondrial division (Lee et al., 2016).

Comparatively little is known about inner mitochondrial membrane (IMM) division, which must also occur for successful mitochondrial division. Early results in *C. elegans* showed that Drp1-deficient animals had an overall mitochondrial division defect but that matrix markers segregated (Labrousse et al., 1999). Drp1-independent mitochondrial constrictions have been observed (Lee and Yoon, 2014), and these constrictions appear to be through direct effects on the IMM (Cho et al., 2017). The constrictions occur at ER-mitochondrial contact sites, depend on increased intra-mitochondrial calcium, and precede full mitochondrial division (Cho et al., 2017).

In this paper, we show that INF2-mediated actin polymerization on ER is necessary for mitochondrial calcium increase upon stimulation with either histamine or ionomycin. This calcium increase requires MCU, which is also required for stimulus-induced IMM contractions and mitochondrial division. INF2-mediated actin polymerization stimulates ER-to-mitochondrial calcium transfer by enhancing close ER-mitochondrial contact. During stimulus-induced mitochondrial division, the IMM divides prior to OMM division.

## Results

### Stimulus-induced actin polymerization enhances mitochondrial Ca^2+^

Studies from our lab and others show that a variety of stimuli causing increased cytosolic calcium, including ionomycin and histamine, trigger a transient cytosolic actin polymerization “burst” (Shao et al., 2015; Ji et al., 2015; Wales et al., 2016). These stimuli operate by distinct mechanisms to raise cytosolic calcium, with ionomycin requiring extracellular calcium while histamine relies only on calcium from intracellular stores (Figure S1A). We previously showed that the ionomycin-induced actin burst stimulates Drp1 oligomerization and subsequent mitochondrial division (Ji et al., 2015). Histamine stimulation also induces Drp1 oligomerization and mitochondrial division, albeit to lower levels than ionomycin (Figure S1B-D).

Since both ionomycin and histamine also induce an increase in mitochondrial matrix calcium (Rizzuto et al., 1993; Abramov and Duchen, 2003), we investigated the relative kinetics of changes in cytoplasmic calcium and mitochondrial calcium in relation to the actin burst using live-cell imaging. Both ionomycin and histamine trigger rapid increases in cytoplasmic calcium, with T_1/2_ of 4.5±1.1 and 3.4±0.4 sec respectively (Figure 1A-F; Video 1,2). The actin burst lags significantly upon ionomycin stimulation (8.3±1.8 sec), but its histamine-stimulated rate is indistinguishable from that of cytoplasmic calcium (3.8±0.9 sec) (Figure 2A-F; Video 1,2). The mitochondrial calcium increase occurs after the actin burst for both stimuli, with T_1/2_ of 15.4±3.4 for ionomycin and 7.5±1.9 sec for histamine ((Figure 2A-F; Video 1,2). This result is confirmed by two-color imaging in the same cell for both ionomycin (Figure 2G, Video 3) and histamine (Figure 2H, Video 4). All responses are transient, returning to near baseline within 200 sec for both stimuli. Further information on ionomycin stimulation is provided in Methods and in Figure S10D.

**Figure 1.**
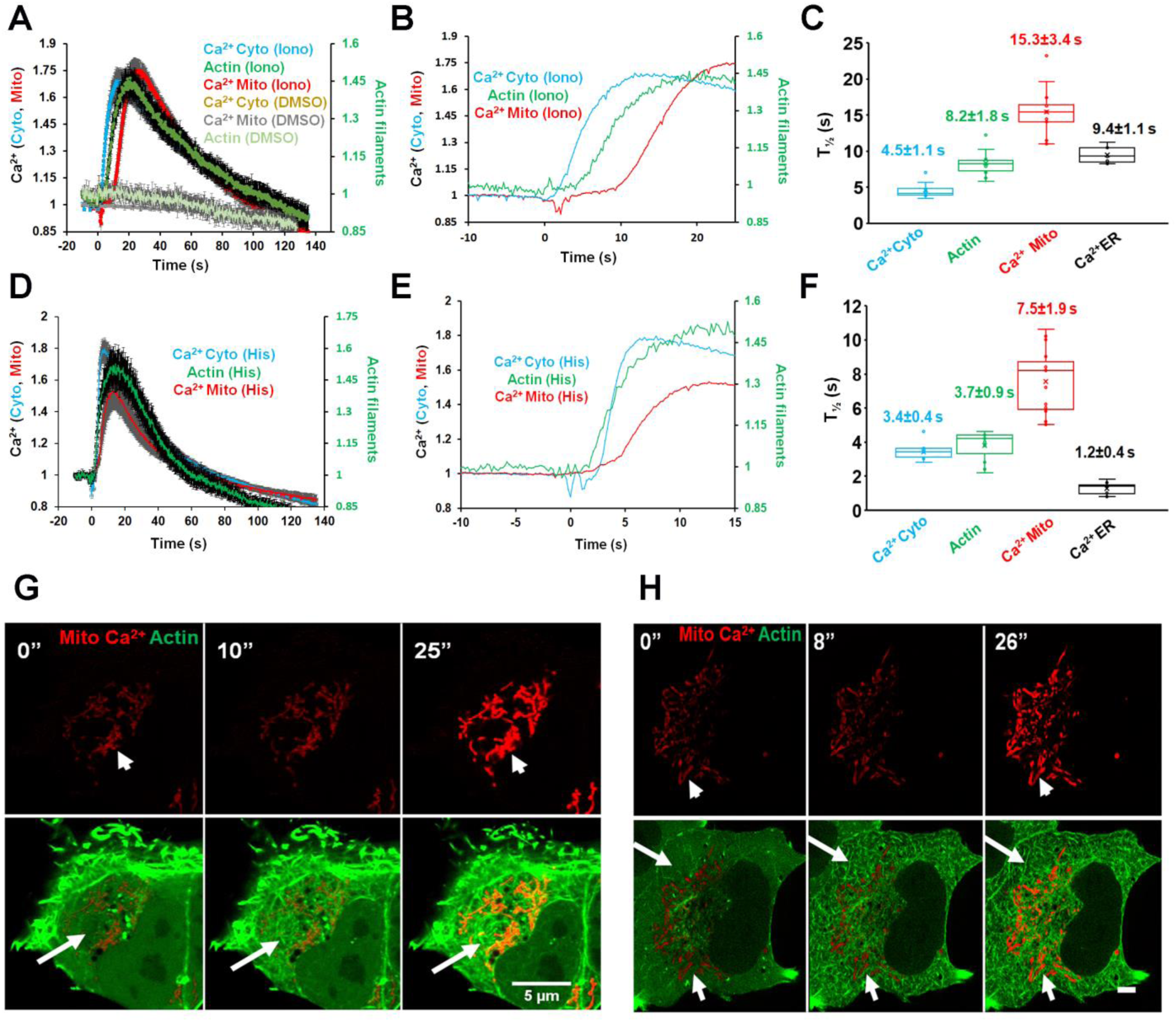
The stimulus-induced actin burst precedes the mitochondrial calcium spike. (A) and (D) Quantification of cytoplasmic calcium (Cyto-R-GECO), cytoplasmic actin (GFP-Ftractin) or mitochondrial calcium (Mito-R-GECO) in U2OS cells after ionomycin (A), 4 μM) or histamine (D), 100 μM) stimulation. Rapid acquisition mode (4.8 frames/sec). Values on Y axis represent the value at time X normalized to the value at time 0 (F/F0). N = 10-16 cells (also given in Figure 1C). Error bars represent standard error of mean (SEM). Corresponds to Videos 1 and 2. (B) and (E) Zoom of early time points from graphs in A (ionomycin) and D (histamine), showing the distinct kinetics of cytoplasmic calcium, actin and mitochondrial calcium. Error bars removed for clarity. (C) and (F) Box-and-whiskers plots of stimulation half-times based on the curves in graphs A and D. Number of cells for each reading is 14 (cyto calcium, ionomycin), 10 (actin, ionomyin), 14 (mitochondrial calcium, ionomycin), 12 (ER calcium, ionomycin), 14 (cyto calcium, histamine), 11 (actin, histamine), 16 (mitochondrial calcium, histamine), 10 (ER calcium, histamine). Each point represents one cell. (G) and (H) Time-lapse montages of actin (green, GFP-Ftractin) and mitochondrial calcium (red, mito-R-GECO) changes in the same cell after ionomycin (G, 4 μM) or histamine (H, 100 μM) stimulation. Mitochondrial calcium alone shown in top panels, and merge in bottom panels. Taken at one frame per 1.03 sec (G) and 1.2 sec (H). Arrow indicates region of the cytoplasm displaying the cytoplasmic actin burst. Corresponds to Videos 3 and 4.

**Figure 2.**
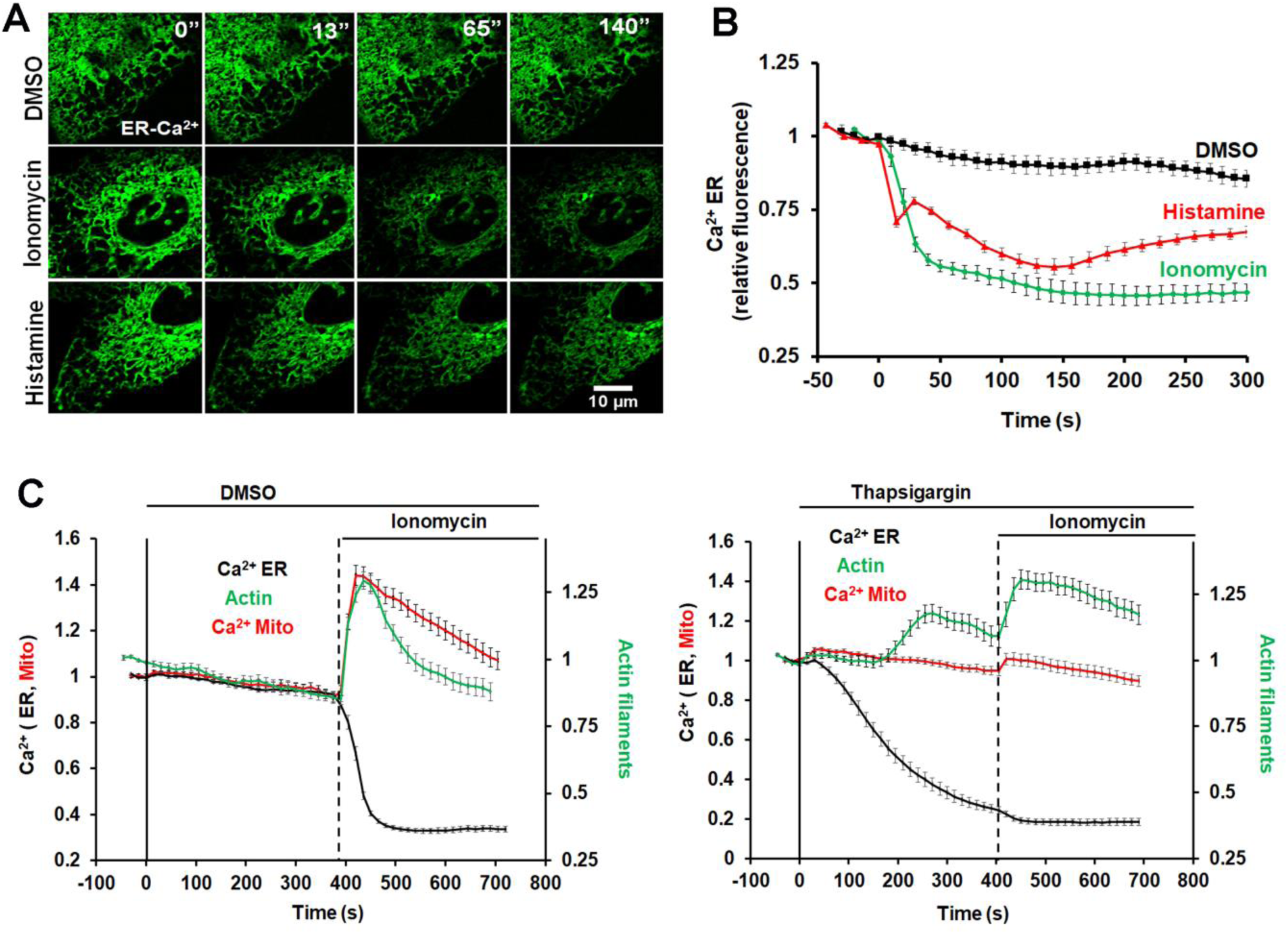
Calcium release from ER triggered by ionomycin and histamine. **A)** Time-lapse montages of U2OS cells transfected with the ER calcium probe (ER-GCamP6-150) and treated with DMSO (top), 4 μM ionomycin (middle) or 100 μM histamine (bottom). Time in sec. Corresponds to Video 5. **B)** Graph quantifying changes in ER calcium upon the treatments described in A. N = 12, 15 and 15 cells for DMSO, ionomycin and histamine, respectively. **C)** Effect of thapsigargin on ER calcium (black curves), actin burst (green curves), and mitochondrial calcium (red curves) before and after ionomycin stimulation. U2OS cells transfected with ER calcium probe and mApple-Ftractin. DMSO (left graph) or thapsigargin (right graph, 1 μM) applied at 0 sec (bold line) and ionomycin (4 μM) applied at the time indicated by the dashed line. N= 10 cells (DMSO); 13 cells (thapsigargin).

We probed in more detail the calcium source for the mitochondrial calcium spike. Since mitochondrial calcium responses often occur at sites of close ER contact (DeStefani et al., 2016), we directly measured changes in ER calcium upon ionomycin or histamine stimulation, using a low-affinity calcium probe for ER, ER-GCaMP6-150 (150 μM K_d_ for Ca^2+^ (de Juan-Sanz et al., 2017)). As shown previously (Montero et al., 1997; Montero et al., 2003), histamine induces a rapid decrease in ER calcium that occurs in two phases (Figure 2A, B; Video 5). The first phase precedes both the actin burst and the overall cytosolic calcium increase (T_1/2_ 1.27 ±0.36 sec, Figure 1F), whereas the second phase occurs 20-60 sec after stimulation. Interestingly, ionomycin also induces a rapid decrease in ER calcium (Figure 2A,B, Video 5) with a T_1/2_ of 9.6 ± 1.1 sec (Figure 1C). This result had been suggested previously but not directly measured (Morgan and Jacob, 2003; Caridha et al., 2008) and implies that a significant component of the ionomycin-induced cytoplasmic calcium increase derives from intracellular stores, likely through calcium-mediated calcium release (Endo, 2009).

We used thapsigargin pre-treatment to test whether ER calcium release was necessary for the mitochondrial calcium increase upon either histamine or ionomycin stimulation. Thapsigargin treatment depletes ER calcium significantly within 10 min (Figure 2C). The resulting rise in cytoplasmic calcium is sufficient to activate actin polymerization, but this activation is significantly slower than the acute activation caused by ionomycin or histamine (Figure 2C), with a T_1/2_ of >200 sec. No mitochondrial calcium increase occurs upon thapsigargin stimulation. For histamine, stimulation after thapsigargin-induced depletion of ER calcium results in no increase in cytoplasmic calcium or mitochondrial calcium, similar to past studies (Diarra and Sauvé, 1992), and histamine does not induce an actin burst (Figure S2A-C). For ionomycin stimulation, thapsigargin pre-treatment reduces the increases in cytoplasmic calcium and actin polymerization, and the increase in mitochondrial calcium is eliminated (Figure 2C, Figure S2D). These results suggest that the mitochondrial calcium increase results largely from ER calcium for both histamine and ionomycin stimulation.

Since the actin burst precedes the mitochondrial calcium spike, we asked whether actin polymerization was necessary for increased mitochondrial calcium. Treatment with the actin sequestering molecule Latrunculin A (LatA) strongly inhibits the mitochondrial calcium spike upon either ionomycin or histamine stimulation (Figure 3A, B), while actually increasing the magnitude of the cytosolic calcium increase (Figure S3A). These results suggest that actin polymerization is required for efficient stimulus-induced mitochondrial calcium entry.

**Fig. 3.**
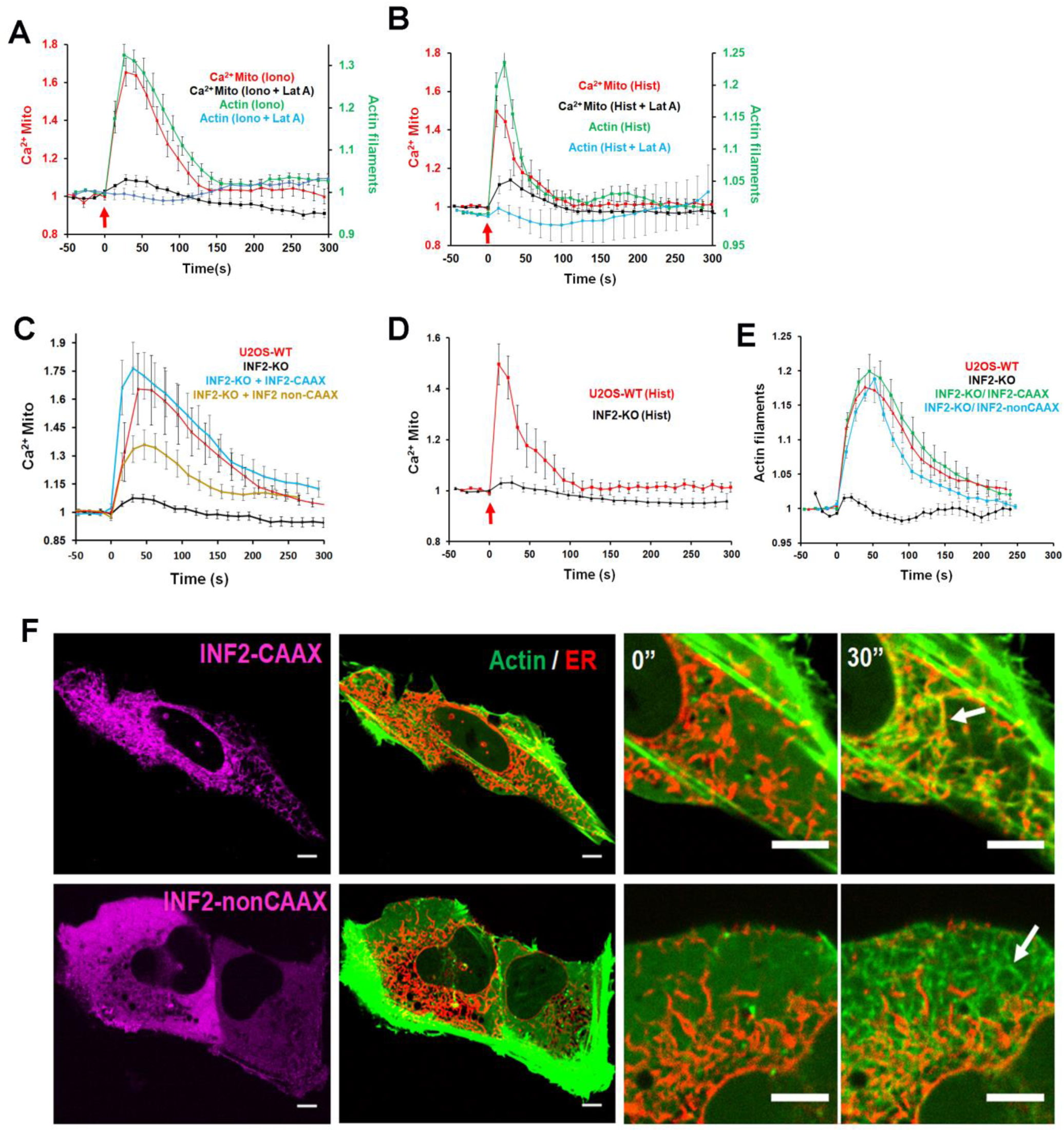
Actin requirement for stimulus-induced mitochondrial calcium spike. A) and B) Effect of Latrunculin A treatment (2 uM) on ionomycin-induced (A) or histamine-induced (B) actin polymerization burst and mitochondrial calcium spike in U2OS cells. N = 10-15 cells. Error bars, SEM. C) and D) Mitochondrial calcium spike following ionomycin (C) or histamine (D) stimulus in U2OS-WT, U2OS-INF2-KO, or INF2-KO cells re-expressing GFP-INF2-CAAX or GFP-INF2-nonCAAX. Control U2OS-INF2-KO cells are expressing GFP-Sec61? as an ER marker instead of INF2. E) Actin polymerization burst following ionomycin in U2OS-WT and U2OS-INF2-KO cells. Rescue by transfection of GFP-INF2-CAAX or GFP-INF2-nonCAAX also shown. N=15-18 cells. Error bars, SEM. F) Ionomycin-induced actin morphology in INF2-KO cells re-expressing GFP-INF2-CAAX or GFP-INF2-nonCAAX. Whole cell view shown pre-stimulation. Insets shown pre-stimulation (0”, left) and after 30 sec stimulation (30”, right). Taken in a single confocal plane in and apical region of the cell body, which reduces actin background from stress fibers but causes the ER to appear fragmented. Scale bar: Main panel: 5 μm; inset: 2 μm. Corresponds to Video 6.

### Actin polymerization on ER enhances ER-to-mitochondrial Ca^2+^ transfer

We previously showed that an ER-bound isoform of the formin INF2, INF2-CAAX, plays a role in mitochondrial division (Korobova et al., 2013) and that siRNA-mediated INF2 suppression eliminates the ionomycin-induced actin burst (Ji et al., 2015). We therefore asked whether INF2-CAAX plays a role in the mitochondrial calcium spike. CRISPR-mediated knock-out of INF2 in U2OS cells (Figure S3B, C) eliminates both the actin burst and the mitochondrial calcium spike upon either ionomycin or histamine stimulation (Figure 3C, D, Figure S3D), while causing an increase in cytoplasmic calcium (Figure S3A). Re-expression of INF2-CAAX or INF2-nonCAAX as GFP fusions restores the actin burst (Figure 3E) but the resulting actin morphology differs between the two isoforms, with INF2-CAAX-induced actin enriching around ER whereas INF2-nonCAAX-induced actin is not ER-enriched (Figure 3F, C, Video 6). Interestingly, INF2-CAAX restores the mitochondrial calcium spike to a greater degree than INF2-nonCAAX (Figure 3C, Figure S4), suggesting that ER localization is important.

These results suggest that INF2-mediated actin polymerization enhances ER-to-mitochondrial calcium transfer. One mechanism to mediate this enhancement is by promoting close contact between ER and mitochondria. Evidence for ER-mitochondrial contact has been obtained previously using low-affinity calcium sensors tethered to the cytoplasmic face of the OMM, showing that increases in ER-mitochondrial contact cause increases in calcium concentration near the OMM that are in excess of the bulk cytoplasmic calcium concentration (Csordás et al., 2010; Giacomello et al., 2010). We used a similar low-affinity calcium probe (LAR-GECO-1.2, K_d_ 12 μM for calcium, Wu et al., 2014) tethered to the cytoplasmic face of the OMM through MAS70. Stimulation of WT U2OS cells by ionomycin causes an increase in OMM-calcium signal, in a similar time course to the mitochondrial calcium increase, and this OMM-calcium increase is greatly attenuated in INF2-KO cells (Figure 4A). Re-expression of INF2-CAAX rescues the ionomycin-stimulated OMM-calcium increase fully, while re-expression of INF2-nonCAAX causes only partial rescue (Figure 4A).

**Figure 4.**
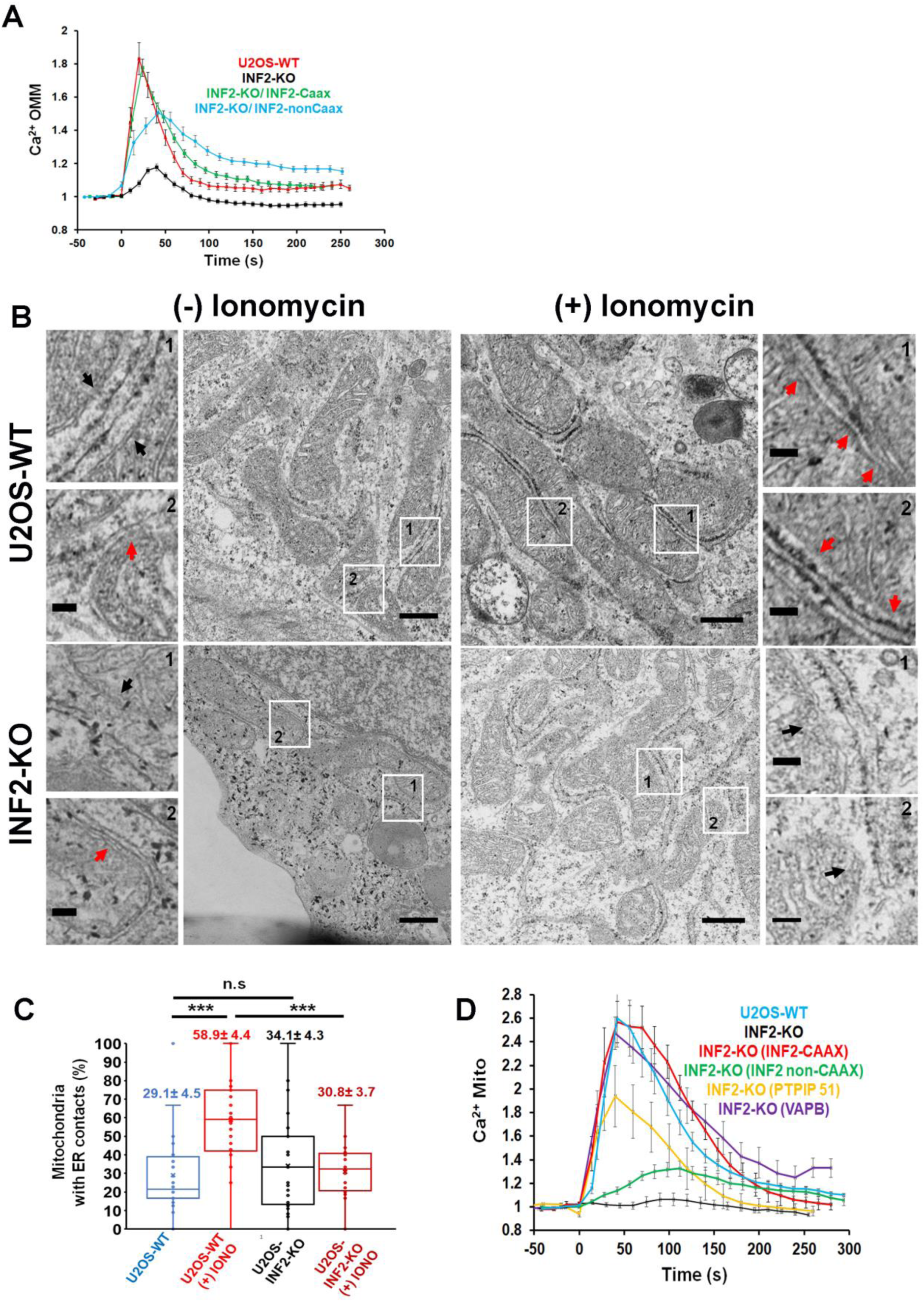
INF2-mediated actin polymerization is required for stimulation of ER-mitochondrial contact. **A)** Change in calcium levels at the cytoplasmic face of the OMM upon ionomycin stimulation (4 μM) in U2OS-WT and INF2-KO cells. Cells were transfected with the Mass70-LAR-GECO1.2 construct, a low-affinity calcium probe (K_d_ 12 μM) tethered to the cytoplasmic face of the OMM. For rescue, INF2-KO cells were transfected with plasmid expressing GFP-INF2-CAAX or GFP-INF2-nonCAAX. Ionomycin added at 0 sec. N= 24 cells (WT); 19 cells (INF2-KO); 22 cells (INF2-CAAX) and 16 cells (INF2-non CAAX). Error bars: SEM **B)** Electron micrographs showing examples of relationship between ER and mitochondria in WT cells (upper images) and INF2-KO cells (lower images) in either the un-stimulated condition ((-) Ionomycin, left images) or after 60 sec stimulation with 4 μM ionomycin ((+) ionomycin, right images). Two zooms shown for each panel (regions indicated by numbered arrows on main panels). Examples of ER-mitochondrial contacts of <30 nm are indicated by red arrowheads, while examples of more distant contacts indicated by black arrowheads. Scale bars represent 500 nm (main micrographs) and 100 nm (zooms). **C)** Quantification from electron micrographs of % mitochondria with close ER contacts in WT cells (N = 244 mitochondria); WT cells stimulated with ionomycin (N = 176 mitochondria), INF2-KO cells (N = 245 mitochondria), and INF2-KO cells stimulated with ionomycin (N = 204 mitochondria). Values represent Mean±SEM; *** p<0.001. Each point represents one imaged field. **D)** Rescue of mitochondrial calcium response in INF2-KO cells by over-expression of ER-mitochondrial tethers. INF2-KO U2OS cells were transfected with a mitochondrial matrix calcium probe (Mito-R-GECO) along with either CFP-VAPB or GFP-PTPIP51, then stimulated with 4 μM ionomycin. The effects of either GFP-INF2-CAAX or GFP-INF2-nonCAAX re-expression are shown for comparison. N= 10 cells (WT); 8 cells (INF2-KO); 11 cells (INF2-nonCAAX); 15 cells (INF2-CAAX); 10 cells (PTPIP51); 14 cells (VAPB). Error bars, SEM.

We also measured changes in ER-mitochondrial contact directly by thin section electron microscopy, defining regions of <30 nm distance as ER-mitochondrial contact sites (Csordás et al., 2006). Prior to ionomycin stimulation, WT and INF2-KO cells contain a similar percentage of mitochondria with close ER contacts (Figure 4B, C). Upon ionomycin stimulation, the percentage of mitochondria with close ER contacts increases 2-fold in WT cells but shows no increase in INF2-KO cells (Figure 4B, C). These results suggest that INF2-mediated actin polymerization increases ER-mitochondrial contact.

We next asked whether over-expression of ER-mitochondrial tethering proteins could compensate for the lack of ionomycin-induced mitochondrial calcium in INF2-KO cells. For this purpose we chose VAPB (on ER) and PTPIP51 (on mitochondria), since over-expression of either of these tether partners individually has been shown to increase ER-mitochondrial tethering (Gomez-Suaga et al., 2017). Indeed, expression of either protein significantly restores the ionomycin-induced mitochondrial calcium spike in INF2 KO cells (Figure 4D, Figure S4). Taken together, these results suggest that actin polymerization by INF2-CAAX enhances ER-mitochondrial contact, leading to increased stimulus-induced mitochondrial calcium.

We have also reported that myosin IIA suppression by siRNA reduces ionomycin-triggered mitochondrial division while not affecting the actin burst (Ji et al., 2015). MyoIIA-KD also attenuates the mitochondrial calcium spike for both ionomycin and histamine stimulation while having similar levels of actin burst in both cases (Figure S5). Taken together, these results suggest that both INF2-mediated actin polymerization and myosin IIA activity are necessary for stimulus-induced mitochondrial calcium entry.

### A role for mitochondrial calcium in mitochondrial division

We next asked whether the mitochondrial calcium spike plays a role in mitochondrial division. MCU is the major IMM channel for mitochondrial calcium, and accounts for most histamine-induced mitochondrial calcium entry (Baughman et al., 2011; De Stefani et al., 2011). In U2OS cells, MCU-targeted siRNA (MCU-KD, Figure S6A) eliminates the ionomycin-induced mitochondrial calcium spike without affecting the cytosolic calcium spike or the magnitude and duration of the actin burst (Figure 5). MCU-KD also decreases baseline fluorescence of mito-R-GECO by 2.5-fold (Figure S6D, E), suggesting a decrease in basal mitochondrial calcium. Re-expression of an siRNA-resistant GFP-MCU construct rescues the ionomycin-induced mitochondrial calcium spike (Figure S6C, D).

**Figure 5.**
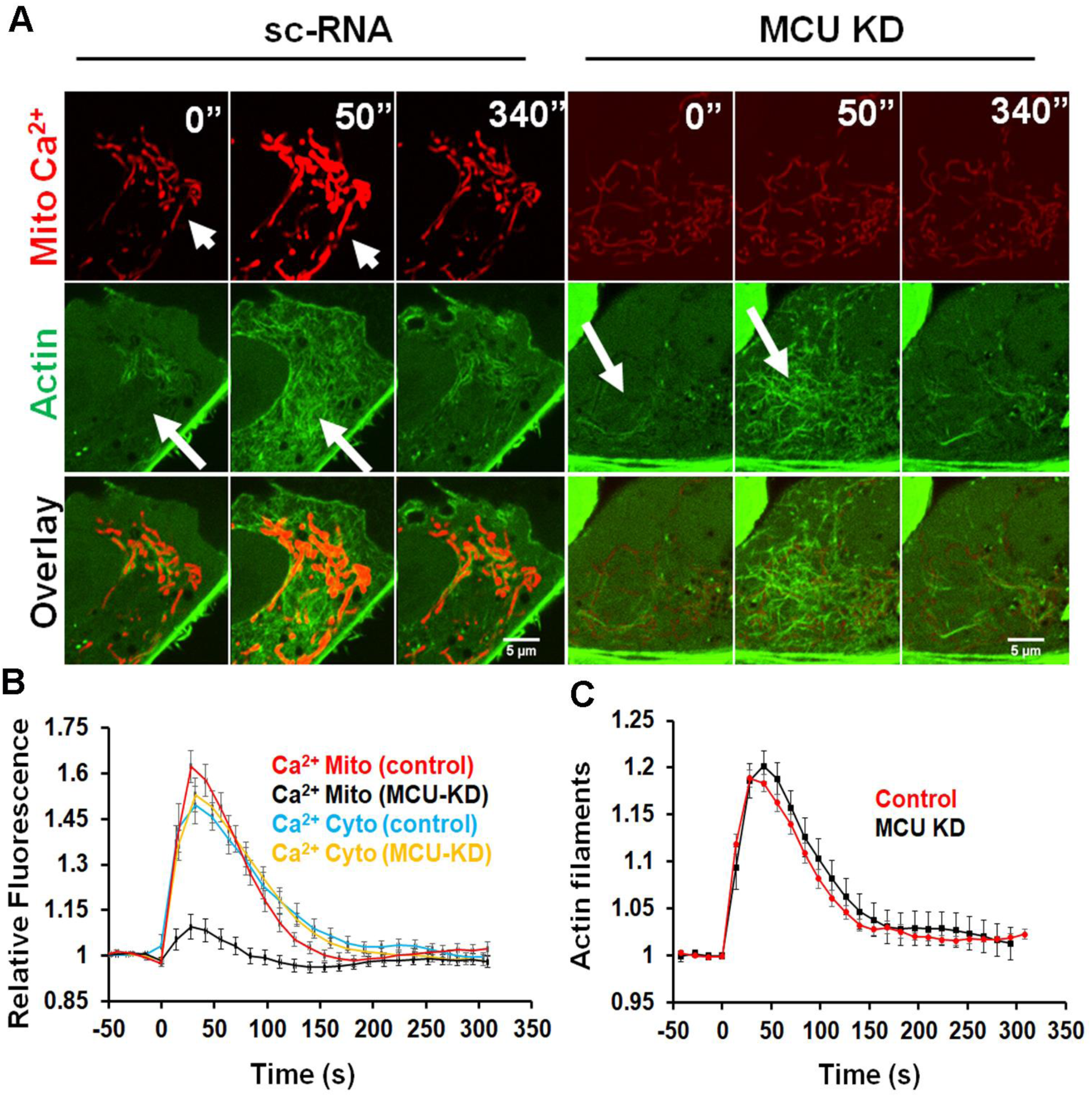
MCU suppression inhibits the mitochondrial calcium spike without inhibiting the actin burst. U2OS cells were transfected with either scrambled siRNA (control) or siRNA against MCU (MCU KD). After 48 hrs, cells were transfected with mito-R-GECO (mitochondrial calcium) and GFP-Ftractin (polymerized actin) or with cyto-R-GECO (cytoplasmic calcium) and imaged at 72 hrs post-siRNA treatment. **A)** Confocal image montages of cells at indicated times after 4 μM ionomycin stimulation. For MCU KD, the mitochondrial calcium signal is enhanced to reveal the faint mitochondrial outline. Arrowheads show mitochondrial calcium rise and arrows denote polymerized actin increases. Scale bar: 5μm. **B)** Quantification of the mitochondrial and cytosolic calcium spikes upon stimulation with 4 μM ionomycin (at time 0). N= 10 cells (Mito calcium control); 18 cells (mito calcium MCU KD); 10 cells (Cyto calcium control); 15 cells (Cyto calcium MCU KD). **C)** Quantification of the actin burst upon stimulation with 4 μM ionomycin (at time 0). N= 10 cells (Control); 15 cells (MCU KD).

We used MCU-KD cells to ask whether mitochondrial calcium plays a role in mitochondrial division. Using a recently established morphometric assay (Lee et al., 2016), MCU-KD causes an increase in the mean area of individual mitochondria, intermediate between control cells and Drp1-KD cells (Figure S7A-C). Given the possibility that the increase in length could be due to either decreased division or increased fusion, we also used a live-cell assay to measure mitochondrial division rate directly. In unstimulated cells, MCU-KD causes a 2.5-fold decrease in division rate (Figure 6A). Ionomycin stimulation induces a 3-fold increase in division rate for WT cells but no significant increase in MCU KD cells (Figure 6A). These results suggest that mitochondrial calcium entry plays a role in mitochondrial division, particularly in division stimulated by elevated calcium.

**Figure 6.**
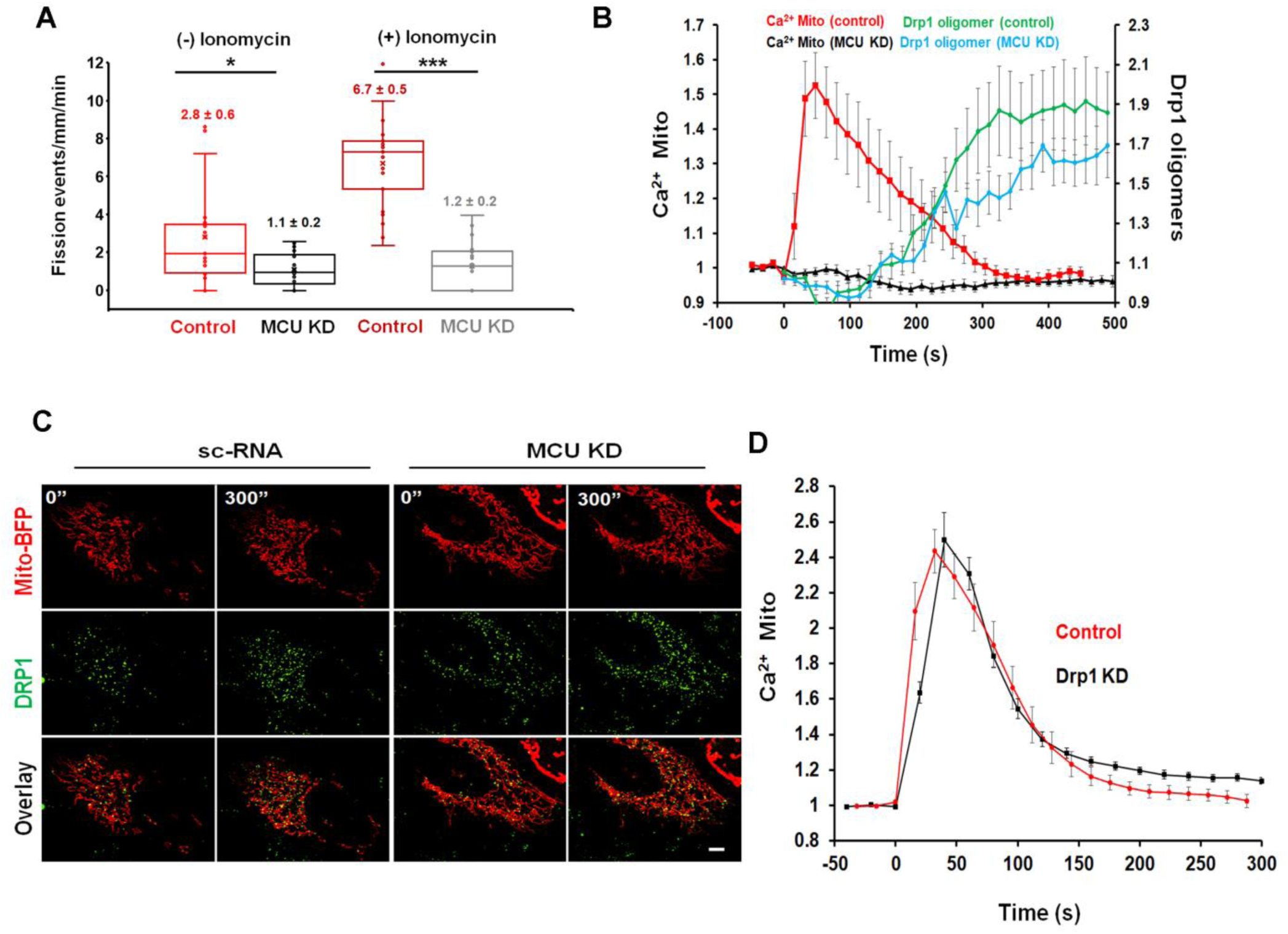
MCU suppression inhibits mitochondrial division but not Drp1 oligomerization. **A)** Quantification of mitochondrial division rate in control and MCU KD U2OS cells by measuring the number of division events in peripheral ROIs from live-cell videos of mitochondrial matrix marker (mito-BFP), either in un-stimulated cells or cells in the first 10 min of ionomycin (4 μM). stimulation. N= 19 cells (Control (-) ionomycin); 21 cells (Control (+) ionomycin); 20 cells (MCU KD (-) ionomycin); 22 cells (MCU KD (+) ionomycin). Each point represents one ROI/ cell. **B)** Quantification of mitochondrially-associated Drp1 oligomers and mitochondrial calcium in response to ionomycin (4 μM). GFP-Drp1 knock-in cells were treated with scrambled siRNA or MCU siRNA for 48 hrs, then transfected with mito-R-GECO and mito-BFP. Cells were imaged at 72 hrs post-siRNA treatment. Ionomycin addition at 0 sec. **C)** Micrographs from GFP-Drp1 knock-in U2OS cells (Control and MCU siRNA treated) transfected with mito-BFP (red) and treated with 4 μM ionomycin as in panel B. Pre-stimulation (0”) and 300 sec stimulations shown. Scale Bar: 5 μm. **D)** Quantification of mitochondrial calcium increase in control or Drp1 siRNA-treated (72 hrs) U2OS cells upon ionomycin (4μM) stimulation. N= 14 cells in each case. Error bars, SEM.

Since Drp1 is a key mitochondrial division factor, and we previously showed that ionomycin causes increased Drp1 oligomerization (Ji et al., 2015), we tested the effect of MCU-KD on Drp1 oligomerization. In a CRISPR-derived GFP-Drp1 knock-in cell line (Ji et al., accepted at *J. Cell Biol*.), MCU suppression does not significantly alter ionomycin-induced Drp1 oligomerization on mitochondria (Figure 6B, C). Since Drp1 can oligomerize on peroxisomes as well as on ER independently of mitochondria (Ji et al., accepted at *J. Cell Biol.*), we also assessed total cellular Drp1 oligomers and found similar ionomycin-induced oligomerization kinetics between control and MCU KD cells (Figure S7D). Conversely, Drp1 suppression does not alter the ionomycin-induced mitochondrial calcium increase (Figure 6D). These results suggest that the role of mitochondrial calcium in mitochondrial division is un-related to Drp1 recruitment.

Interestingly, siRNA of either Drp1 or MCU causes increased expression of the other protein, with MCU KD causing a 2-fold increase in Drp1, and Drp1 KD causing a 2-fold increase in MCU (Figure S6A, B). Furthermore, siRNA of INF2 causes ∼2-fold increases in both Drp1 and MCU (Figure S6A, B). These results may suggest compensatory changes to maintain mitochondrial division upon depletion of key components.

### Drp1-independent IMM constriction

Mitochondria in un-stimulated U2OS cells periodically constrict simultaneously along their length (Figure S8A), as reported previously for other cell types (Lee and Yoon, 2014; Cho et al., 2017). Only rarely do these constrictions undergo division. Interestingly, constrictions accompany a transient rise in matrix calcium (Figure S8B). The distance between constrictions is approximately 2 μm (Figure S8C).

Ionomycin causes a 6.7-fold increase in mitochondrial constrictions, on a similar time scale to the mitochondrial calcium increase (Figure 7A, B, Video 7). By time-lapse Airyscan microscopy, 90.4 ± 1.9% of the constrictions correspond to sites of close ER association (Figure 7C, D, Video 8). We used siRNA suppression to ask whether constrictions were due to Drp1-induced OMM constriction. Surprisingly, Drp1-KD causes an increase in mitochondrial constrictions for both the unstimulated and ionomycin-stimulated conditions (Figure 7A, B, Video 8). These constrictions can be so exaggerated that aberrant bulges appear at adjacent regions. We next used MCU-KD cells to test whether increased mitochondrial calcium was necessary for these constrictions. MCU-KD results in a significantly reduced constriction number for both the unstimulated and ionomycin-stimulated conditions (Figure 7A, B, Video 7). These findings suggest that the mitochondrial constrictions result from mitochondrial calcium influx, not Drp1 activity.

**Figure 7.**
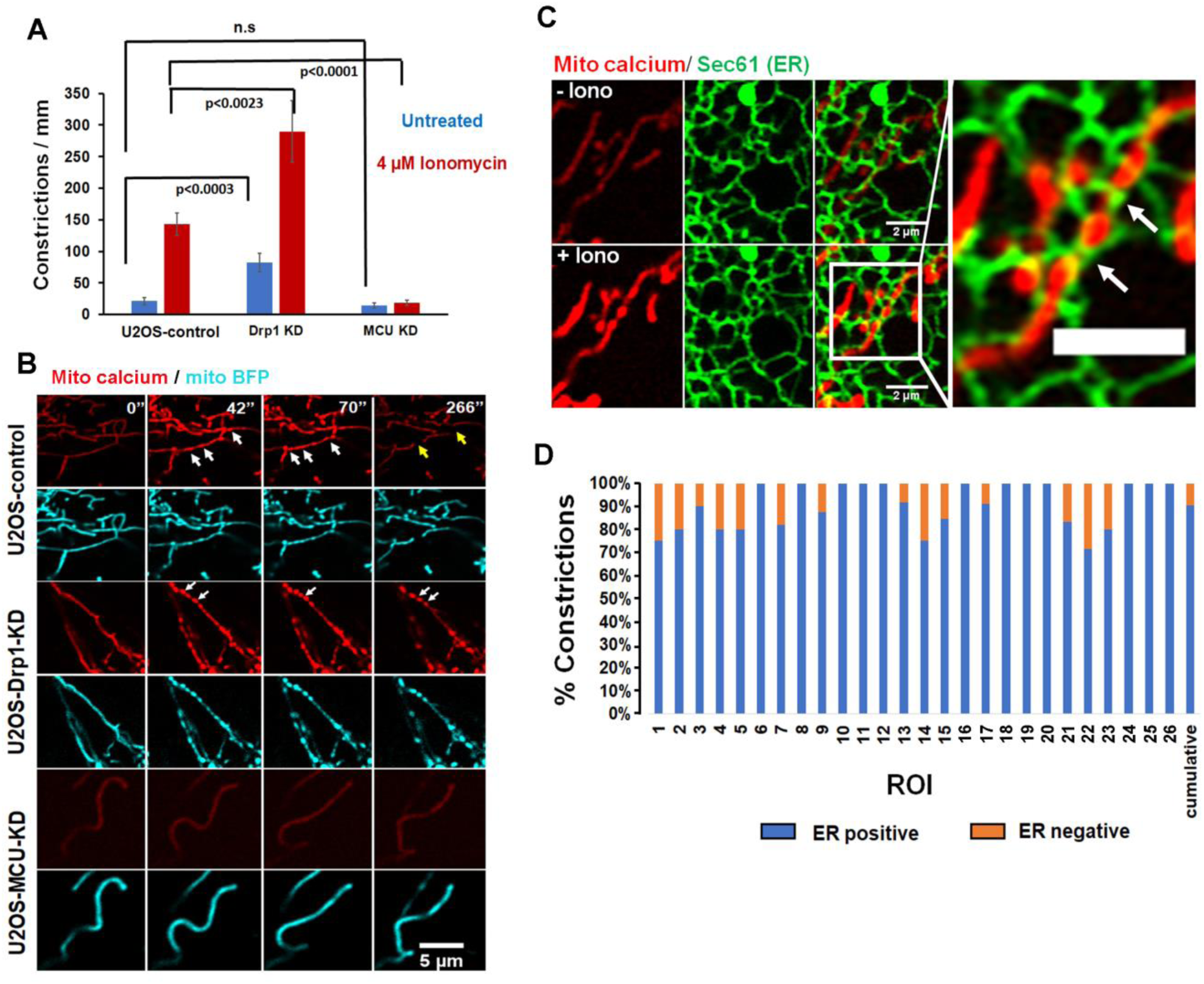
Mitochondrial constrictions are MCU-dependent and Drp1-independent. **A)** Quantification of mitochondrial constriction frequency in the unstimulated (blue) or ionomycin-stimulated (red) conditions, represented as constrictions per mm mitochondrial length in U2OS-control, Drp1-KD and MCU-KD cells. Error bars, SEM. Control: 16 ROI, 2508 μm mitochondrial length; Drp1 KD: 10 ROI, 1000 μm; MCU KD: 20 ROI, 2600 μm. **B)** Representative confocal montages of control, Drp1-KD and MCU-KD U2OS cells transfected with mito-R-GECO (mitochondrial calcium, red) and mitoBFP (mitochondrial matrix, blue), then treated with ionomycin (4 μM) at time 0. Time in sec. Arrows: white-constrictions; yellow-fission event. Corresponds to Video 7. **C)** U2OS cell transfected with GFP-Sec61? (green, ER marker) and mito-R-GECO (red, mitochondrial calcium) and imaged live before ionomycin stimulation (top) and at 35 sec after 4 μM ionomycin addition (bottom). Image at right is expanded view of + ionomycin condition. Scale bar: 2 μm in regular panel; 1μm in expanded panel. Corresponds to Video 8. **D)** Quantification of percent mitochondrial constrictions corresponding to ER contact from live-cell confocal microscopy of ionomycin-stimulated cells such as in Figure 7C/Video 9. 204 constrictions from 26 ROIs/13 cells assessed, totalling 461 μm^2^ mitochondrial area.

Our results suggest that mitochondrial constrictions are due to IMM dynamics. To examine dynamics of both IMM and OMM, we used Airyscan microscopy of cells expressing mito-R-GECO and GFP-Tom20 (an OMM marker). During ionomycin-stimulated mitochondrial division events, a consistently observable intermediate stage exists in which no discernable matrix marker is visible while the OMM marker presents as two parallel lines (Figure 8A, B, Video 9, and Figure S9A). Similar results occur on a comparatively slower time scale upon treatment with CGP37157 (Figure S9B, C, Video 10), an inhibitor of the mitochondrial Na^+^/Ca^+^ anti-porter, a major route of mitochondrial calcium exit (Kaddour-Djebbar et al., 2010; Chowdhury et al., 2011). From the CGP37157 data, the mean lag time between apparent matrix separation and apparent OMM separation for 29 measured events is 32.3 ± 29.1 sec (29 events, Figure 8C).

**Figure 8.**
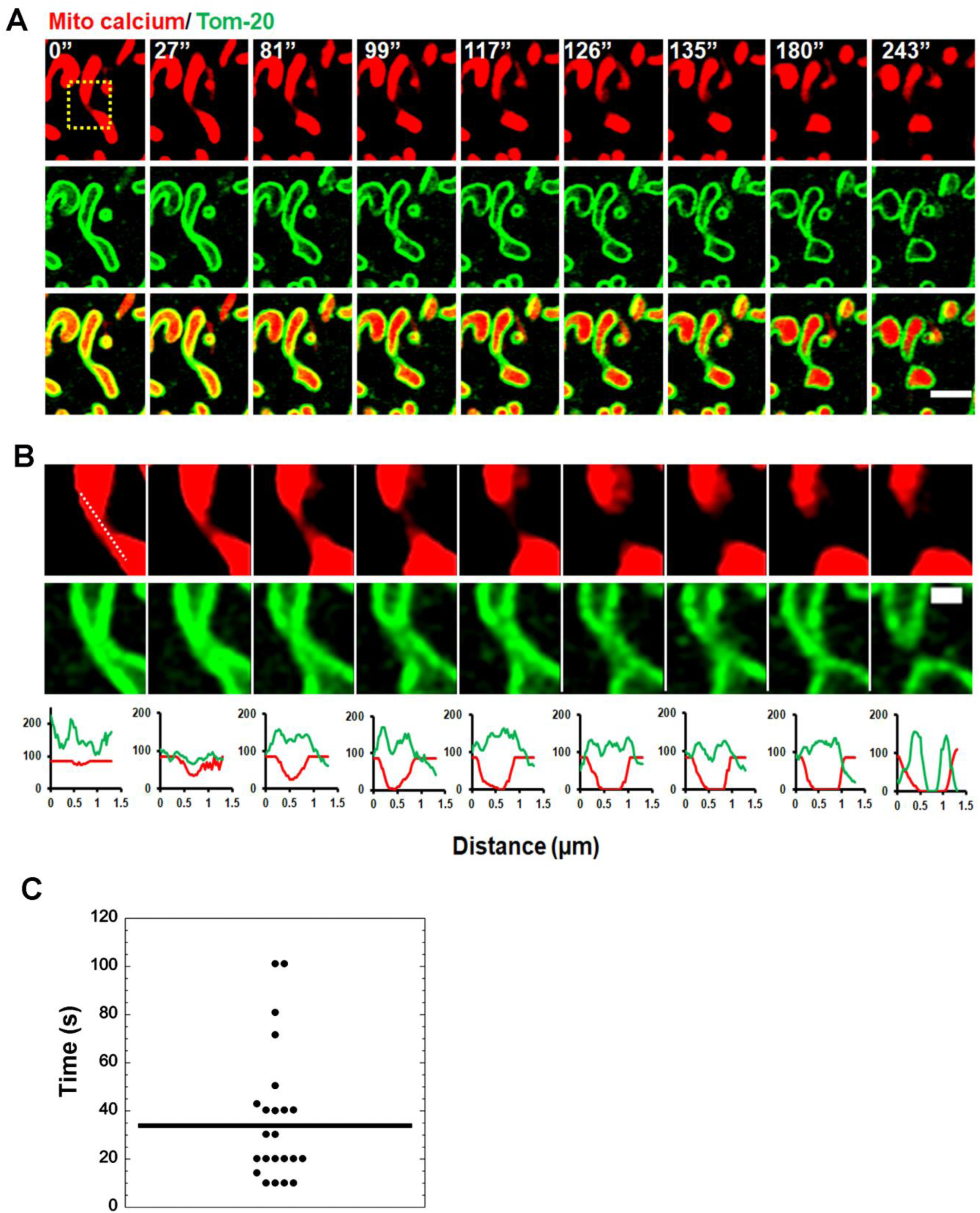
IMM division prior to OMM division during mitochondrial division. **A)** U2OS cells transfected with mito-R-GECO (mitochondrial matrix calcium, red) and GFP-Tom20 (OMM, green) were imaged live by Airyscan microscopy at 9 sec intervals after ionomycin treatment (4 μM). Representative time-lapse montage of a mitochondrial division event shown here. Scale bar: 2 μm. Correspond to Video 9 **B)** Zoom of the division site (boxed in panel A). The mito-R-GECO levels has been enhanced in an attempt to detect the existence of a thin matrix tether. Line scans indicate that the OMM tether persists in the absence of a matrix tether. Scale bar: 500 nm. **C)** Quantification of time between apparent matrix separation and apparent OMM separation in mitochondrial division events after CGP37157 stimulation (example in Figure S9B). 29 division events from 23 cells. Mean time, 32.3±29.1 sec. Error bars, standard deviation. Each point represents one division event.

We probed the mechanism by which increased matrix calcium leads to IMM constriction by testing the role for Opa1, a dynamin GTPase that is attached to the IMM. Opa1 is well known to mediate IMM fusion, but targeted proteolysis releases Opa1 from the IMM and may allow Opa1 to participate in IMM division (Anand et al., 2014; MacVicar & Langer, 2016). One study shows evidence that Opa1 is not required for constrictions (Lee and Yoon, 2014) while another study shows that Oma1 processing of Opa1 is necessary for constrictions (Cho et al., 2017). We probed for changes in Opa1 proteolytic processing upon ionomycin stimulation and found no apparent change during the time period of increased IMM constrictions (Figure 9A). Using a covalent cross-linking assay adapted from others (Otera et al., 2016; Figure S10A), there is no apparent change in Opa1 oligomerization upon ionomycin stimulation (Figure 9B). In addition, ionomycin-induced constrictions are still present in U2OS cells upon Oma1 suppression (Figure 9C), despite effectively blocking Opa1 proteolytic processing (Figure S10B, C). These results suggest that Opa1 processing by Oma1 is not required to induce IMM constrictions.

**Figure 9.**
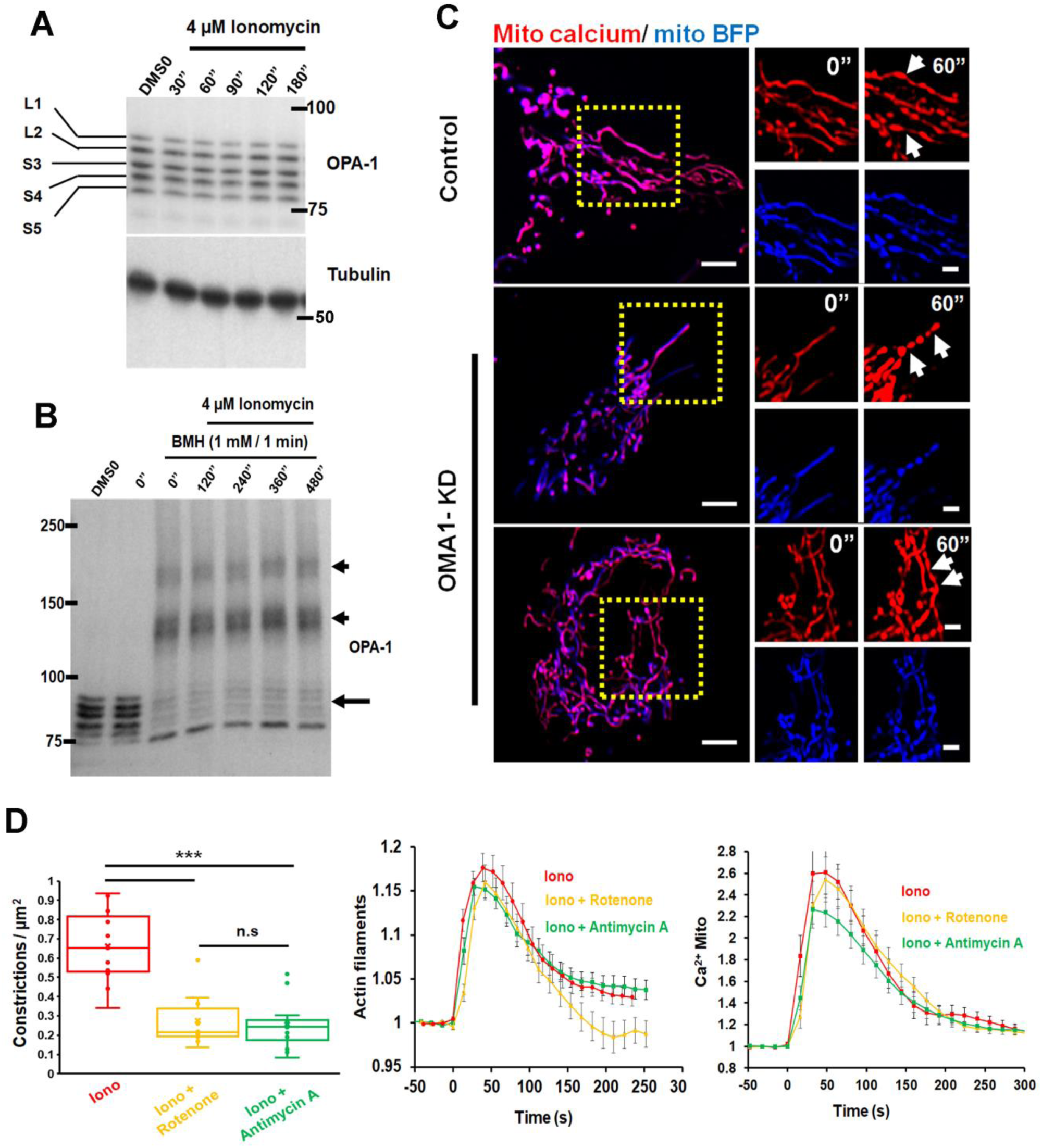
IMM constrictions do not require Oma1 activity but do require electron transport chain. **A)** Western blot of Opa1 isoform pattern in U2OS cells during ionomycin stimulation (4 μM) during the period of optimum calcium-induced constrictions. **B)** Western blot of Opa1 after live-cell crosslinking with BMH, to reveal Opa1 oligomerization changes during ionomycin stimulation (4 μM). Oligomers are labeled with arrowheads and monomers with arrow. **C)** Constrictions in U2OS cells that had been transfected with Oma1 siRNA for 72 hrs, followed by transfection of mito-R-GECO and mitoBFP for 24 hr and stimulation by ionomycin (4 μM) for 60 sec. Scale bar: 5 μm (Main Panel); 2μm (Inset). **D)** Effect of antimycin A (5 μM) or rotenone (2.5 μM) on (left to right): mitochondrial constrictions, actin burst, and mitochondrial calcium spike induced by ionomycin (4 μM). Antimycin A or rotenone was added simultaneously to ionomycin. For mitochondrial constriction assessment: N= 10 ROI, 200μm^2^ mitochondrial area (Iono); 11 ROI, 136 μm^2^ (Iono + Rotenone); 16 ROI, 300 μm^2^ (Iono + Antimycin A). For, actin filament assessment, 10 cells for each condition. For mitochondrial calcium assessment, 15-20 cells for each condition.

We also asked whether a calcium-mediated increase in electron transport chain (ETC) activity was necessary for IMM constrictions, since calcium activates dehydrogenases associated with the ETC. Treatment with rotenone (complex I inhibitor) or antimycin A (complex III inhibitor) significantly decreases ionomycin-induced mitochondrial constrictions, without decreasing the actin burst or mitochondrial calcium spike (Figure 9D). Since the ETC inhibitors are added simultaneous to the ionomycin stimulation, these results suggest that a calcium-mediated increase in ETC activity is required for IMM constrictions.

## Discussion

The combination of work here and in past studies (Korobova et al., 2013, Ji et al., 2015) demonstrates that INF2-mediated actin polymerization on ER stimulates mitochondrial division by two independent mechanisms: (i) mitochondrial calcium uptake, leading to IMM constriction and (ii) Drp1 oligomerization, leading to OMM constriction. Actin stimulates the mitochondrial calcium increase by enhancing ER-mitochondrial interaction (this study), allowing more efficient flow of calcium from ER to mitochondrion. Independently, actin stimulates Drp1 oligomerization through direct binding (Ji et al., 2015; Hatch et al., 2016). Mitochondrial calcium uptake is rapid, with a T_1/2_ of <20 sec, and triggers constriction of the IMM in a manner requiring the electron transport chain. Drp1 oligomerization occurs on a slower time course, with a T_1/2_ of ∼100 sec.

INF2 is activated by increased cytoplasmic calcium (Shao et al., 2015; Wales et al., 2016). We postulate that an initial rise in cytoplasmic calcium induced (stimulated by ionomycin or histamine here) causes INF2 activation, with the resulting ER-localized actin polymerization driving ER-mitochondrial contact, which then allows efficient calcium transfer to mitochondria through MCU. The relative kinetics of cytoplasmic calcium, actin polymerization and mitochondrial calcium in response to these stimuli support this mechanism.

This is the first study demonstrating a role for actin polymerization in ER-to-mitochondrial calcium transfer. A past study showed that 3-hr treatment with actin depolymerizing drugs reduced ER calcium release from hippocampal neurons (Wang et al., 2002), but the length of treatment raised the possibility of indirect effects. Here, we show that ionomycin- or histamine-stimulated mitochondrial calcium increase is blocked by LatA applied simultaneously with the stimulus, or by INF2 knock-out. These stimuli activate calcium dynamics by two distinct mechanisms. Histamine activates ER calcium efflux through IP3 receptors (Berridge et al., 2003). Ionomycin causes initial calcium entry from the extracellular medium, which activates a secondary release of calcium from the ER. This “calcium-induced calcium release” from the ER is presumably largely through ryanodine receptors (Smith et al., 1986). While ryanodine receptors are best characterized in muscle or neuronal systems (Endo, 2009), they are also widely expressed in non-excitable cells (Querfurth et al., 1998; Giannini et al., 1995; Bennett et al., 1996; Park et al, 2014). It is known that a disproportionate amount of ER-released calcium enters the mitochondrion, instead of dissipating into the bulk cytosol. Close contacts of < 30 nm between ER and mitochondrion are thought to allow preferential mitochondrial calcium entry (Rizzuto et al 1998; Csordás et al., 2006; Giacomello et al., 2010; Csordás et al., 2010), and a number of proteins that mediate ER-mitochondrial tethering might be important in this interaction (Phillips and Voeltz, 2016).

It is not clear what population of actin filaments enhances ER-mitochondrial interaction. The INF2-mediated actin “burst” occurs throughout the cytosol (Shao et al., 2015; Ji et al., 2015; Wales et al., 2016), but we have shown that the ER-bound INF2 isoform is specifically associated with mitochondrial dyanmics, and show here and elsewhere (Ramabhadran et al., 2012) that this isoform stimulates actin polymerization on ER. One model is that INF2 assembles actin directly at ER-mitochondrial contacts, perhaps in association with mitochondrially-bound Spire1C (Manor et al., 2015). Short filamentous structures have been observed at ER-mitochondrial close contacts by thin section electron microscopy (Csordás et al., 2006), but their molecular identity is unknown. The fact that even relatively stable actin-based structures are destroyed during traditional electron microscopy processing (Lehrer, 1981; Maupin and Pollard, 1983) suggests that the short actin filaments required to span ER-mitochondrial contacts (10-20 subunits) are likely to be largely eliminated during EM investigation of these contacts. Another model is that actin exerts its effect on ER-mitochondrial contact from a distance, perhaps by decreasing motility of each organelle, as we have observed for mitochondria (Korobova et al., 2013). In addition to actin polymerization directly at mitochondrial division sites, we have observed polymerization parallel to the mitochondrion (Ji et al., 2015), which could serve such a purpose.

Regardless of the mechanism, other molecules contribute to actin-mediated enhancement of ER-mitochondrial contact. The fact that myosin IIA is required for the mitochondrial calcium increase suggests that it is providing contractile tension for ER-mitochondrial contact. The aforementioned Spire1C has been shown to work with INF2 in other aspects of mitochondrial division (Manor et al., 2015). Other studies have shown evidence for actin “clouds” assembling around mitochondria, and these clouds correspond to increased mitochondrial division (Li et al., 2014; Moore et al., 2016). It is unclear whether these structures constitute the same process as that mediated by INF2, since the morphology of the actin filaments differs significantly.

In light of the bulk of our results, it is intriguing that thapsigargin treatment successfully elicits a cytoplasmic actin burst, but fails to stimulate an increase in mitochondrial calcium. One difference is that the thapsigargin-induced actin burst is significantly slower than those elicited by ionomycin or histamine, the former requiring ∼100 sec while the latter peak within 10 sec. This difference in kinetics may speak to the specific actin pool that must be activated to enhance ER-mitochondrial contact, rather than the bulk cytoplasmic actin pool. Alternately, activation of other molecules, such as myosin IIA through phosphorylation (Heissler and Sellers, 2016), might be insufficient in the slower thapsigargin response. More detailed examination of specific actin and myosin populations should shed light on this question.

The demonstration that IMM constricts prior to OMM division during mitochondrial division, both here and in a recently published study (Cho et al., 2017), suggests that specific mechanisms for IMM constriction must exist. In this study, we find that constrictions are strongly suppressed by electron transport chain inhibitors (antimycin, rotenone). We apply these inhibitors simultaneous to the constriction stimulus, suggesting that an acute metabolic increase is necessary. Increased matrix calcium is known to activate pyruvate dehydrogenase and isocitrate dehydrogenase (Denton, 2009), linking the actin-stimulated mitochondrial calcium rise to IMM constrictions through increased metabolic flux.

At present, the mechanism by which increased metabolic flux drives IMM constriction is unclear. Cristae remodeling is an attractive possibility. A major player in cristae remodeling is the dynamin GTPase Opa1, which is bound to the IMM and mediates mitochondrial fusion, but might also mediate IMM division upon proteolytic processing that releases it from the IMM (Anand et al., 2014, MacVicar and Langer, 2016). Opa1 is also involved in the maintenance of cristae morphology, as well as regulating OMM/IMM tethering through the MICOS complex (Cogliati et al., 2016; Otera et al., 2016; Quintana-Cabrera et al., 2017). Findings from two studies differ on the importance of Opa1 to mitochondrial constrictions. One study (Lee and Yoon, 2014) shows that spontaneous mitochondrial constrictions do not require Opa1, but that Opa1 is required for subsequent IMM proton leak. Another study (Cho et al., 2017) shows that Oma1 or Opa1 suppression inhibits IMM constrictions in cortical neuron cultures. Possible explanations for these differences include distinct constriction mechanisms based on cell type or stimulus. A second Opa1-processing protease, Yme1 (Anand et al., 2014, MacVicar and Langer, 2016) has not yet been tested for its importance in constrictions and, while Oma1 is the relevant protease in neurons (Cho et al., 2017), there may be redundancy in other cells.

We show for the first time that apparent IMM division precedes OMM division. We use the term “apparent” because it is possible that a thin matrix tether persists that is undetectable in our studies, even though over-processing of the matrix signal fails to detect such a tether. Despite this caveat, we observe a period of tens of seconds in which there is no detectable matrix and the OMM appears parallel at the eventual division site, strongly suggestive of IMM division prior to OMM division. However, we have never observed stable IMM division in the absence of eventual OMM division, suggesting that the two events are coupled. A repeated observation in the field has been that a Drp1-independent “pre-constriction” occurs prior to full mitochondrial division (Labrousse et al., 1999; Legesse-Miller et al., 2003). Calcium-stimulated IMM constriction/division may be the origin of such pre-constriction.

## Acknowledgements

We thank Tom Blanpied, Sai Divakaruni, Minerva Contreras, Mike Hoppa, Stefan Strack, Charles Barlowe, Bill Wickner, Anna Hatch, Mu A, Lorna Young, Laura McCormick, Lori Schoenfeld, and Yuriy Usachev for advice, reagents and preliminary experiments that strengthened this work, A. Lavanway for frequent microscopy help, and Uma Clic for the ability to rise when asked. Supported by NIH GM069818, GM106000 and DK88826 to HNH and NIH NS036942 to TAR. Purchase of the Airyscan microscope was possible through generous support from NIGMS (Supplement to GM109965), the Provost and Dean of Sciences at Dartmouth College, and the Norris Cotton Cancer Center. Purchase of Dragonfly microscope was possible through generous support from the William Ruger Senior Fund at Dartmouth (560118) and the Department of Biochemistry and Cell Biology.

## Author Contributions

R.C. and H.N.H. conceived of or designed the research and wrote the manuscript. R.C., R. S. and W-K.J. performed the experiments. J.J.S. and T.A.R. designed ER Ca^2+^ experiments.

## Methods

### Plasmids and siRNA oligonucleotides

Mito-R-GECO1, Cyto-R-GECO1 (K_d_= 0.48 μM for calcium), and Mito-LAR-GECO1.2 (K_d_= 12 μM for calcium) constructs were kind gifts from Yuriy M. Usachev (Dept of Pharmacology, University of Iowa Carver College of Medicine) and have been described previously (Wu et al., 2014). The OMM-calcium probe was a kind gift from Stefan Strack (Dept. of Pharmacology, University of Iowa Carver College of Medicine) and was constructed from the Mito-LAR-GECO1.2 plasmid by replacing the matrix targeting sequence with the OMM targeting sequence from *S. cerevisiae* Mas70 (amino acids 1-25). mApple-F-tractin and GFP-F-tractin plasmid was a gift from Clare Waterman and Ana Pasapera (NIH, Bethesda, MD), and described in (Johnson and Schell, 2009). Mito-DsRed and mito-BFP constructs were previously described (Korobova et al., 2014), and consist of amino acids 1–22 of S. cerevisiae COX4 N-terminal to the respective fusion protein. ER-tagRFP was a gift from Erik Snapp (Albert Einstein College of Medicine, New York City), with prolactin signal sequence at 5’ of the fluorescent protein and KDEL sequence at 3’. GFP-hSec61β was a kind gift from Prof. Tom Rapoport (Harvard Medical School, Boston). Tom20-GFP was made by restriction digest of Tom20 from Tom20-mCherry (kind gift from Andrew York, NIH) with NheI and BamHI and cloning into eGFP-N1 (Clonetech). ER-GCaMP6-150 (K_d_= 150 uM for calcium) was described in (de Juan-Sanz et al., 2017). Mammalian expression vectors for CFP tagged VAPB and eGFP tagged PTPIP51 were a kind gift from Christopher C.J. Miller (King’s College, London) (Gomez-Suaga et al., 2016). The pDEST47-GFP-MCU plasmid for rescue experiments was from Addgene (31732) and described in (Baughman et al., 2011)). The GFP-INF2-CAAX and nonCAAX plasmids used for rescue were similar to those reported in (Ramabhadran et. al., 2013).

Oligonucleotides for all siRNA used were synthesized by IDT and are listed below:

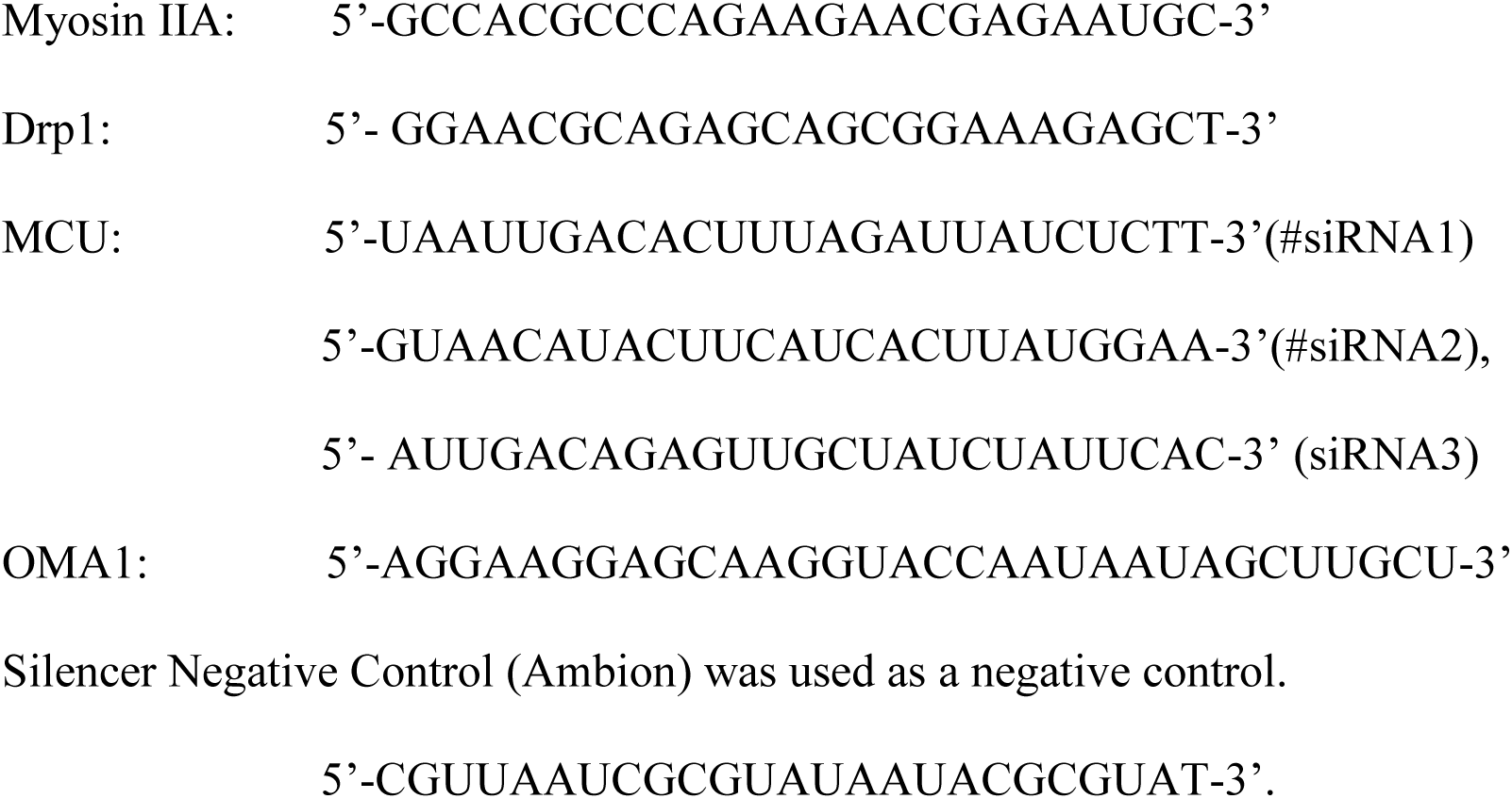

### Antibodies and Western Blotting

Anti-Myosin IIA (Cell Signaling Tech, 3403S) was used at 1:1000. Anti-MCU (D2Z3B Rabbit Ab) (Cell Signaling Tech, #14997) was used at 1:1000. Anti-Drp1 (Cell Signaling Tech, 8570) was used at 1:1000. Anti-GAPDH (Santa Cruz G-9) was used at 1:1500. Anti-Tubulin (DM1-α, Sigma/Aldrich) was used at 1: 10,000 dilutions. Anti-actin (Millipore, mab1501R) used at 1:1000. Anti-INF2 was described previously (Ramabhadran et al., 2011; 941-1249 antibody used). Anti-Opa1 (BD Biosciences, 616606) was used at 1:2000. Anti-Mff (Proteintech, 17090-1-AP) used at 1:1000. Anti-Fis1 (Proteintech, 10956-1-AP) used at 1:1000. Anti-Mid51 (Proteintech) used at 1:500. Anti-OMA1 (Santa Cruz Biotech, sc-515788) used at 1:1000. Anti-Tom20 (Santa Cruz Biotech, sc-145) used at 1:500 for immunofluorescence. Anti-Rabbit Texas Red secondary antibody (Vector Lab, Tl1000) used at 1:500 for Immunofluorescence. For Western blotting, cells were grown on 6 well plate, trypsinized, washed with PBS and resuspended 50 μL PBS. This solution was mixed with 34 μL of 10% SDS and 1 μL of 1 M DTT, boiled 5 minutes, cooled to 23°C, then 17 μl of 300 mM of freshly made NEM in water was added. Just before SDS-PAGE, the protein sample was mixed 1:1 with 2xDB (250 mM Tris-HCl pH 6.8, 2 mM EDTA, 20% glycerol, 0.8% SDS, 0.02% bromophenol blue, 1000 mM NaCl, 4 M urea). Proteins were separated by 7.5% SDS-PAGE and transferred to a PVDF membrane (polyvinylidine difluoride membrane, Millipore). The membrane was blocked with TBS-T (20 mM Tris-HCl, pH 7.6, 136 mM NaCl, and 0.1% Tween-20) containing 3% BSA (Research Organics) for 1 hour, then incubated with the primary antibody solution at 4°C overnight. After washing with TBS-T, the membrane was incubated with horseradish peroxidase (HRP)-conjugated secondary antibody (Bio-Rad) for 1 hour at room temperature. Signals were detected by Chemiluminescence (Pierce).

### Cell culture, transfection and drug treatments

Human osteosarcoma U2OS cells (American Type Culture Collection HTB96) were grown in DMEM (Invitrogen) supplemented with 10% calf serum (Atlanta Biologicals). The GFP-Drp1 KI U2OS cell line made by CRISPR-Cas9 is described elsewhere (W. K. Ji et al, manuscript under review). Briefly, we used the GeCKO system (Zhang laboratory, MIT, http://genome-engineering.org/gecko/). The donor plasmid contained eGFP (A206K mutant) flanked by 445 bases upstream of hDrp1 start codon and 308 bases downstream from start (synthesized by IDT). The target guide sequence (CATTCATTGCCGTGGCCGGC) was predicted using the GeCKO website program and made by IDT. Donor and guide plasmids were transfected into U2OS cells at a 3:1 molar ratio using Lipofectamine 2000 (Invitrogen). Cells were put under puromycin selection and clones were selected by FACS sorting and single cell cloning, then verified by IF and Western blotting. The cell line expresses GFP-Drp1 at 50% of the total Drp1 level, with the remaining Drp1 being un-modified (overall Drp1 level unchanged from control cells).

To prepare the INF2-KO-U2OS cell line, the guide sequence to INF2 (TCCGTGGGGTCCGAATCCTG) was predicted and cloned into LentiCRISPRv2 vector by following the protocol found at http://genome-engineering.org/gecko/ The resulting guide plasmid, along with helper plasmids psPAX2 and pMD2.G, were transfected into HEK293 cells using Lipofectamine LTX (Invitrogen). Media containing transfection reagent was aspirated the next day and replaced with fresh media. Supernatant containing lentivirus was collected 48 hours post-transfection and filtered through a 0.45um filter. Fresh virus was used to infect U2OS cells. After 48 hours, media containing virus was removed and replaced with fresh media containing 2μg/ml puromycin. Cells were put under puromycin selection and clones were selected by FACS sorting into 96 well plates, then verified by immunofluorescence and Western blotting.

For transfection, cells were seeded at 4x10^5^ cells per well in a 6-well dish ∼16 hours prior to transfection. Plasmid transfections were performed in OPTI-MEM media (Invitrogen) with 2 μL Lipofectamine 2000 (Invitrogen) per well for 6 hours, followed by trypsinization and re-plating onto concanavalin A (ConA, Sigma/Aldrich, Cat. No. C5275)-coated glass bottom MatTek dishes (P35G-1.5-14-C) at ∼3.5x10^5^ cells per well (coverslips treated for ∼2 hours with 100g/mL ConA in water at room temperature). Cells were imaged in live cell media (Life Technologies, Cat.No. 21063-029), ∼16-24 hours after transfection.

For all experiments, the following amounts of DNA were transfected per well (individually or combined for co-transfection): 500 ng for Mito-R-GECO1, Cyto-R-GECO1, Mito-LAR-GECO1.2, mt-DsRed, GFP-F-Tractin, mito-BFP; 600 ng for GFP-Tom20; 750 ng for ER-GCaMP6-150; 1000 ng for ER-tagRFP, GFP-Sec61β; 600 ng for CFP-VAPB and GFP-PTPIP51; 500 ng for OMM-Mito calcium construct.

For siRNA transfections, 1x10^5^ cells were plated on 6-well plates, and 2 μl RNAimax (Invitrogen) and 63 pg of siRNA were used per well. Cells were analyzed 72-84-hour post-transfection for suppression.

For ionomycin (Sigma Aldrich-I0634) or histamine (Sigma Aldrich H7125), cells were treated with 4 μM ionomycin (from a 1 mM stock in DMSO) or 100 μM histamine (from 100 mM stock in DMSO) at the fifth frame (about 1 min, depending on time interval used) during imaging and continued for another 5-10 min. Medium was pre-equilibrated for temp and CO2 content before use. DMSO was used as the negative control. Of note, the ionomycin treatment used here differs markedly from those in most other studies in that we conduct our stimulation in the presence of serum. This treatment results in the transient changes in cytoplasmic and mitochondrial calcium shown, with both returning to baseline in 120 sec. In contrast, most other studies treat in the absence of serum. In our hands, serum removal causes a higher and prolonged mitochondrial calcium increase, in addition to extensive fragmentation of both mitochondria and ER (Figure S10D).

For Latrunculin A (LatA, Calbiochem 428021) coupled with ionomycin treatment, cells were treated with live cell medium containing LatA (2 μM) (added from a 2 mM DMSO stock) and ionomycin simultaneously at frame 5 (about 1 min, depends on time intervals) with DMSO used as the negative control. For CGP17357 (Sigma Aldrich Cat No. C8874), cells were treated with 80 μM (from 10 mM stock in DMSO) at frame 10 (about 1 min depending on the time intervals used) during imaging. For Thapsigargin (Life technologies, T7458), cells were treated with 1 μM (from 1 mM stock in DMSO) at frame 4 (about 1 min depending on the time interval used) during imaging. ETC inhibitors (Antimycin A (Sigma A8674)-5 μM final, Rotenone (Sigma 557368)-2.5 μM final) were added simultaneously with ionomycin after 4 frames of imaging (1 min approximate depending on frame speed used).

### Live imaging by confocal and Airyscan microscopy

MatTek dishes were imaged on a Wave FX spinning disk confocal microscope (Quorum Technologies, Inc., Guelph, Canada, on a Nikon Eclipse Ti microscope), equipped with Hamamatsu ImageM EM CCD cameras and Bionomic Controller (20/20 Technology, Inc) temperature-controlled stage set to 37°C. After equilibrating to temperature for 10 min, cells were imaged with the 60x 1.4 NA Plan Apo objective (Nikon) using the 403 nm laser and 450/50 filter for BFP, 491 nm and 525/20 for GFP, 561 nm and 593/40 for RFP. For high time resolution imaging, a Dragonfly 302 spinning disk confocal (Andor Technology Inc, Belfast UK) on a Nikon Ti-E base and equipped with an iXon Ultra 888 EMCCD camera, a Zyla 4.2 Mpixel sCMOS camera, and a Tokai Hit stage-top incubator. Lasers: solid state 405 smart diode 100 mW, solid state 488 OPSL smart laser 50 mW, solid state 560 OPSL smart laser 50 mW, solid state 637 OPSL smart laser 140 mW. Objective: 60x 1.4 NA CFI Plan Apo (Nikon).

Airyscan images were acquired on LSM 880 equipped with 63x/1.4 NA plan Apochromat oil objective, using the Airyscan detectors (Carl Zeiss Microscopy, Thornwood, NY). The Airyscan uses a 32-channel array of GaAsP detectors configured as 0.2 Airy Units per channel to collect the data that is subsequently processed using the Zen2 software. After equilibrating to 37 °C for 30 min, cells were imaged with the 405 nm laser and 450/30 filter for BFP, 488 nm and 525/30 for GFP, 561 nm and 595/25 for RFP.

### Measurements of calcium changes and actin burst in live cells

For the following experiments, imaging was conducted at either 0.21 sec intervals (high time resolution) or 12-15 sec intervals (low time resolution), as indicated. All imaging was in live cell media (DMEM + 4.5 g/l glucose, L glutamine, HEPES-Life technologies [21063-029]) containing 10% Fetal Calf Serum.

#### Calcium measurements

Cells were seeded at 4x10^5^ confluency and transfected with the indicated mitochondrial /cytosolic GECO probes or the ERGCaMP6-150 probe (ER Calcium) prior to the day of imaging. 24 hours after transfections, these cells were imaged for 1 min to establish baseline fluorescence (F0) at 561 nm excitation. Cells were then perfused with the indicated compounds by addition of equal volume of a 2x stock in medium with continuous imaging (F-each time point). Mean fluorescence was calculated for each cell using ImageJ (NIH). Fluorescence values for each time point after drug treatment (F) were normalized with the average initial fluorescence (first 5-6 frames-F0) and plotted against time as F/F0. The rapid mito-R-GECO response is not due to a change in matrix volume, since the fluorescence intensity of co-transfected mitoBFP does not change appreciably.

#### Actin “burst” measurements

Cells were seeded at 4x10^5^ confluency and transfected with GFP-Ftractin prior to the day of imaging. 24 hours after transfections these cells were imaged (at a vertical plane above the basal surface, to avoid actin stress fibers) for 1 min to establish baseline fluorescence at 561 nm excitation. Cells were then perfused with the indicated drugs with continuous imaging. Mean Fluorescence values for each time point were calculated from 4 ROI/cell selected in the peri-nuclear region using ImageJ. Fluorescence values for each time point after drug treatment (F) were normalized with the average initial fluorescence (first 5-6 frames-F0) and plotted against time as F/F0.

#### Mitochondrially associated Drp1 puncta

Drp1-KI cells transiently transfected with mitochondrial markers were imaged live by spinning disc confocal fluorescence microscopy for 10 min at 3 sec intervals in a single focal plane. Regions of interest with readily resolvable mitochondria and Drp1 were processed as described previously (Ji et al., 2015). We thresholded mitochondrially associated Drp1 puncta by using the Colocalization ImageJ plugin with the following parameters: Ratio 50%(0-100%); Threshold channel 1: 30 (0-255); Threshold channel 2: 30 (0-255); Display value: 255 (0-255). Mitochondrially associated Drp1 puncta were further analyzed by Trackmate as described previously (Ji et al., 2015) to separate into low threshold and high threshold categories. The number of Drp1 puncta in each category were automatically counted frame-by-frame using the Find Stack Maxima ImageJ macro.

#### Morphological analysis of mitochondrial size

Following methods described previously (Lee et al., 2016), and adapted for U2OS cells. Briefly U2OS-WT cells (plated on MatTek dishes) were treated with siRNAs against MCU or Drp1 for 72 hrs. The cells were then fixed with 1% Glutaraldehyde in PBS for 10 min and treated with sodium borohydride in PBS to quench auto-fluorescence. These cells were then stained with Tom 20 antibody (mitochondria), anti-rabbit secondary antibody and DAPI (nucleus) in PBS + 10% calf serum, and imaged in PBS on the MatTek dish to prevent flattening due to mounting. Z-stacks (20x0.4 μm) were acquired to obtain the total cell volume. Images were analyzed in a blinded manner (coded by one investigator and analyzed by a second investigator). Maximum intensity projections were generated from z-stacks acquired and background subtracted using FIJI (rolling ball 20.0). 225 μm^2^ area of resolvable mitochondria were selected (one ROI/cell in the spread region of the cell) and analyzed using “Analyze Particle” plugin in FIJI to obtain number of mitochondrial fragments and the area of each fragment per ROI. Images were then de-coded to generate the respective graphs.

#### Mitochondrial division rate

Described in detail previously (Ji et al., 2015). Suitable ROI’s were selected for analysis based on whether individual mitochondria were resolvable and did not leave the focal plane. One RO1 per cell selected. Files of these ROIs were assembled, then coded and scrambled by one investigator, and analyzed for division by a second investigator in a blinded manner as to the treatment condition. The second investigator scanned the ROIs frame-by-frame manually for division events, and determined mitochondrial length within the ROI using the ImageJ macro, Mitochondrial Morphology. The results were then given back to the first investigator for de-coding.

#### Mitochondrial constriction quantification

Confocal time courses were collected of U2OS cells (control, Drp1-KD or MCU-KD) transfected with mitoBFP and mito-R-GECO, recording at 12 sec intervals for 1 min prior to ionomycin stimulation, then for 300 sec after ionomycin stimulation (4 μM in presence of serum). Suitable ROIs were selected for analysis based on whether individual mitochondria were resolvable and did not leave the focal plane (one ROI per cell). Constrictions were scored visually as near-diffraction limited reduction in mitoBFP signal within mitochondria. For each ROI, constrictions were scored for each of the 5 frames prior to stimulation and averaged to give the unstimulated value, and for 10 consecutive frames surrounding the 2-min mark post-stimulation to give the ionomycin-induced value. Number of ROIs and the total length of mitochondria scanned for each set given in the figure legend.

#### Electron Microscopy and analysis of ER-mitochondrial contact

U2OS cells (control, IFN2-KO) were seeded at 40,000 cells/cm^2^ onto glass bottom MatTek dishes and cultured overnight in full growth medium. Next day, the cells were either left untreated or stimulated for 1min with 4 μM ionomycin. Cells were fixed (1h at room temp, two changes of fixative) in 2% glutaraldehyde, 3% paraformaldehyde in 0.1M sodium cacodylate pH7.2; postfixed (1h on ice, in dark) in 1% OsO4 in 0.1M sodium cacodylate pH 7.2; rinsed (2x10min at room temp) in 0.1M sodium cacodylate pH 7.2; incubated (overnight, RT in dark) in 2% aqueous uranyl acetate (EMS, Hatfield, PA), dehydrated in graded ethanol (50%,70%, 95, 2x100%) and embedded in LX-112 resin (Ladd Research, Burlington, VT) using standard protocols (Stan et al., 2012). Parallel sections to the cellular monolayer were obtained using a Leica Ultracut 6 ultramicrotome, the sections mounted on carbon / formvar coated 300 mesh copper grids (Ted Pella, Redding, CA), stained with uranyl acetate and lead citrate (EMS, Hatfield, PA), examined under a Jeol 1010 electron microscope and imaged using a bottom mounted AMT CCD camera (2048x2048 pixels). TIFF images were acquired by one investigator (R. S.) scrambled by another investigator (H.N.H.) and analyzed by a third investigator (R. C.) using FIJI. Mitochondria and ER were located based on their respective morphology and were traced by hand. Distance between the organelles was measured using the ‘measure’ tool in Fiji along the traced-out contact region. Mitochondria with ER within 30 nm at any point on the mitochondrial circumference were scored as a percentage of total mitochondria in the field. The values were de-coded by the second investigator and the graphs generated.

#### Opa1 crosslinking

Crosslinking assay using BMH was modified from the protocols described previously (Otera et al., 2016; Patten et al., 2014). The first paper used a low concentration (50 μM) for 30 min and the second paper used high concentration (1 mM) for 20 min. We first tested both concentrations for varying time (Figure S10A) and decided to use 1 mM BMH for 1 min, in order to accommodate the rapid changes induced by ionomycin.

U2OS cells were treated with 4 μM ionomycin for the indicated times. At 1 min prior to termination, 2X BMH solution (2 mM) in medium was added. The medium was quickly aspirated and replaced with modified SDS-PAGE extraction buffer (250 mM Tris-HCl pH 6.8, 2 mM EDTA, 20% glycerol, 0.8% SDS, 0.02% bromophenol blue, 1000 mM NaCl, 4 M urea) to extract protein for western blotting.

## Supplemental Figure Legends

**Figure S1.**
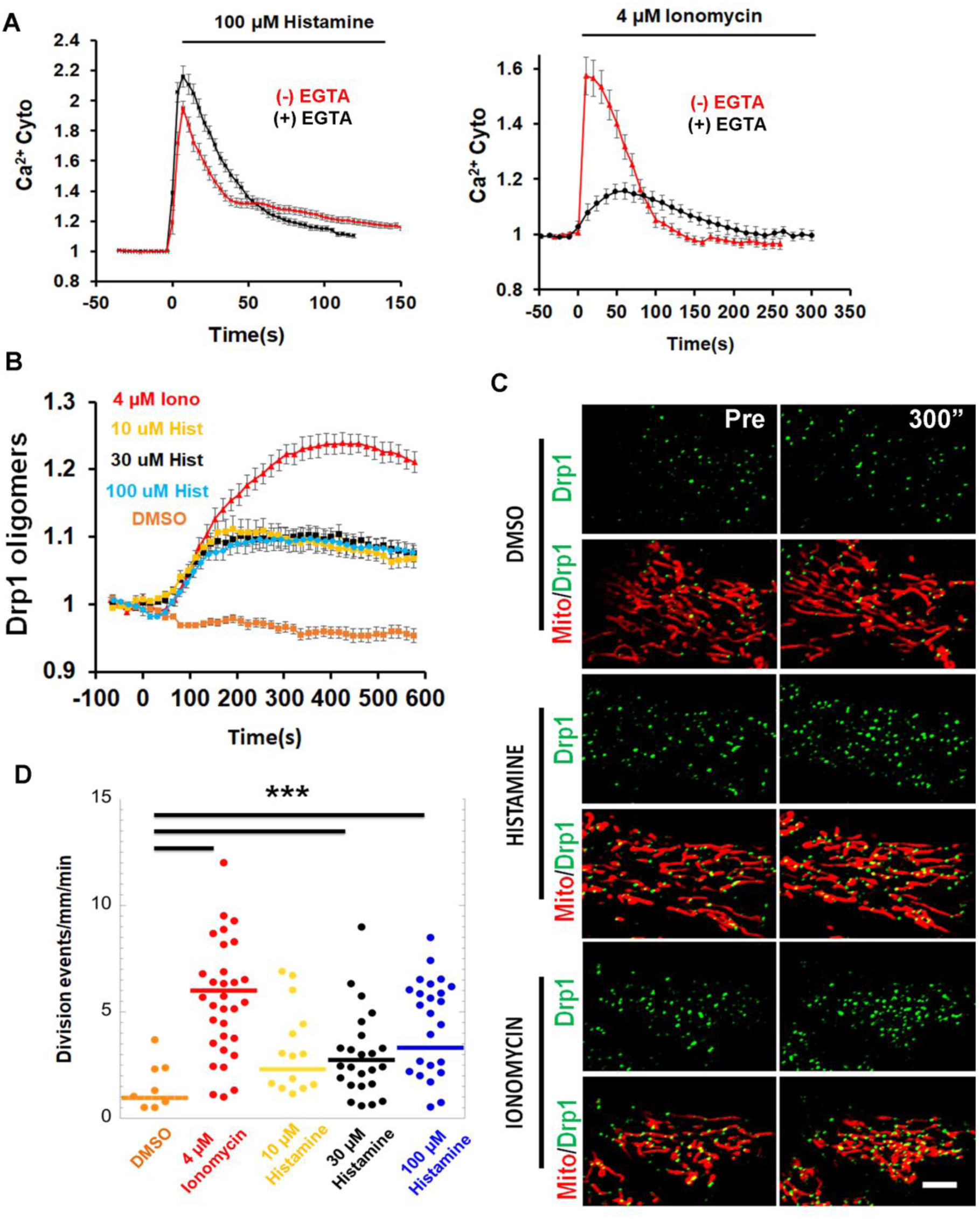
Comparing responses to histamine and ionomycin for cytoplasmic calcium, Drp1 oligomerization, and mitochondrial division. All experiments conducted in continuous presence of 10% calf serum. A) Cytoplasmic calcium. U2OS cells (transfected with Cyto-R-Geco) were stimulated with 100 μM histamine (left) or 4 μM ionomycin (right) in the absence (red) or presence (black) of 5 mM EGTA. N = 12 cells (his); 15 cells (hist + EGTA); 14 cells (Iono); 18 cells (Iono + EGTA). B) Drp1 oligomerization. Quantification of total Drp1 oligomers (whole cell) in response to ionomycin (4 μM) or varying concentrations of histamine (10 μM (yellow); 30 μM (black); 100 μM (blue)) in U2OS cells (GFP-Drp1 knock-in cells transfected with mito-BFP). N= 10-15 cells. C) Micrographs of ROIs from GFP-Drp1 knock-in U2OS cells transfected with mito-BFP (red) and treated with DMSO, 100 μM histamine, or 4 μM ionomycin as in panel B. Left, pre-stimulation. Right, 300 sec stimulation Scale Bar: 5 μm. D) Mitochondrial division rate upon treatment with ionomycin (4 μM) or varying concentrations of histamine (10, 30, or 100 μM) in U2OS cells. DMSO: 13 regions of interest (ROI). Ionomycin- or histamine-treated: at least 20 ROI. Rates (division events/millimeter/min): DMSO: 0.9±1.1; 4 μM Ionomycin: 5.9±3.7; 10 μM Histamine: 2.3±2.2; 30 μM Histamine: 2.7±2.0; 100 μM Histamine: 3.3±2.7.

**Figure S2.**
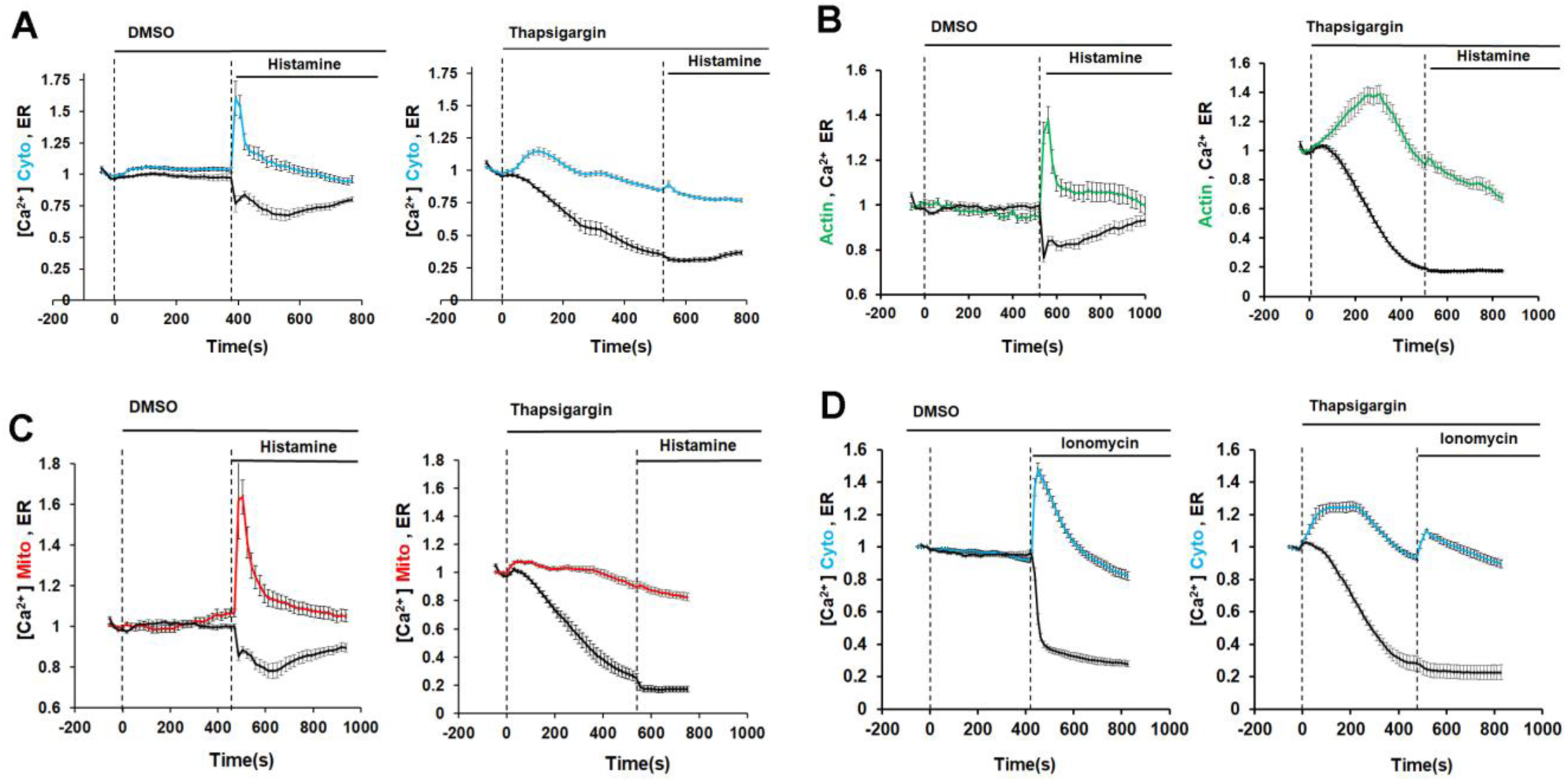
Effects of Thapsigargin on histamine- and ionomycin-induced stimuli. For all panels, DMSO (left graph) or Thapsigargin (1 μM, right graph) applied at 0 sec (first dashed line) and histamine (100 μM) or ionomycin (4 μM) applied at the time indicated by second dashed line. A) Histamine-induced cytoplasmic calcium and ER calcium changes. U2OS cells transfected with ER calcium probe and Cyto-R-GECO. N= 15 cells (DMSO); 15 cells (Thapsigargin). B) Histamine-induced actin burst and ER calcium. U2OS cells transfected with ER calcium probe and mApple-Ftractin. N= 10 cells (DMSO); 11 cells (Thapsigargin). C: Histamine-induced mitochondrial calcium and ER calcium. U2OS cells transfected with ER calcium probe and mito-R-GECO. N= 12 cells (DMSO); 14 cells (Thapsigargin) D: Ionomycin-induced cytoplasmic calcium and ER calcium. U2OS cells transfected with both the Cyto-R-Geco and ER-GCaMP6-150. N= 17 cells (DMSO); 16 cells (Thapsigargin).

**Figure S3.**
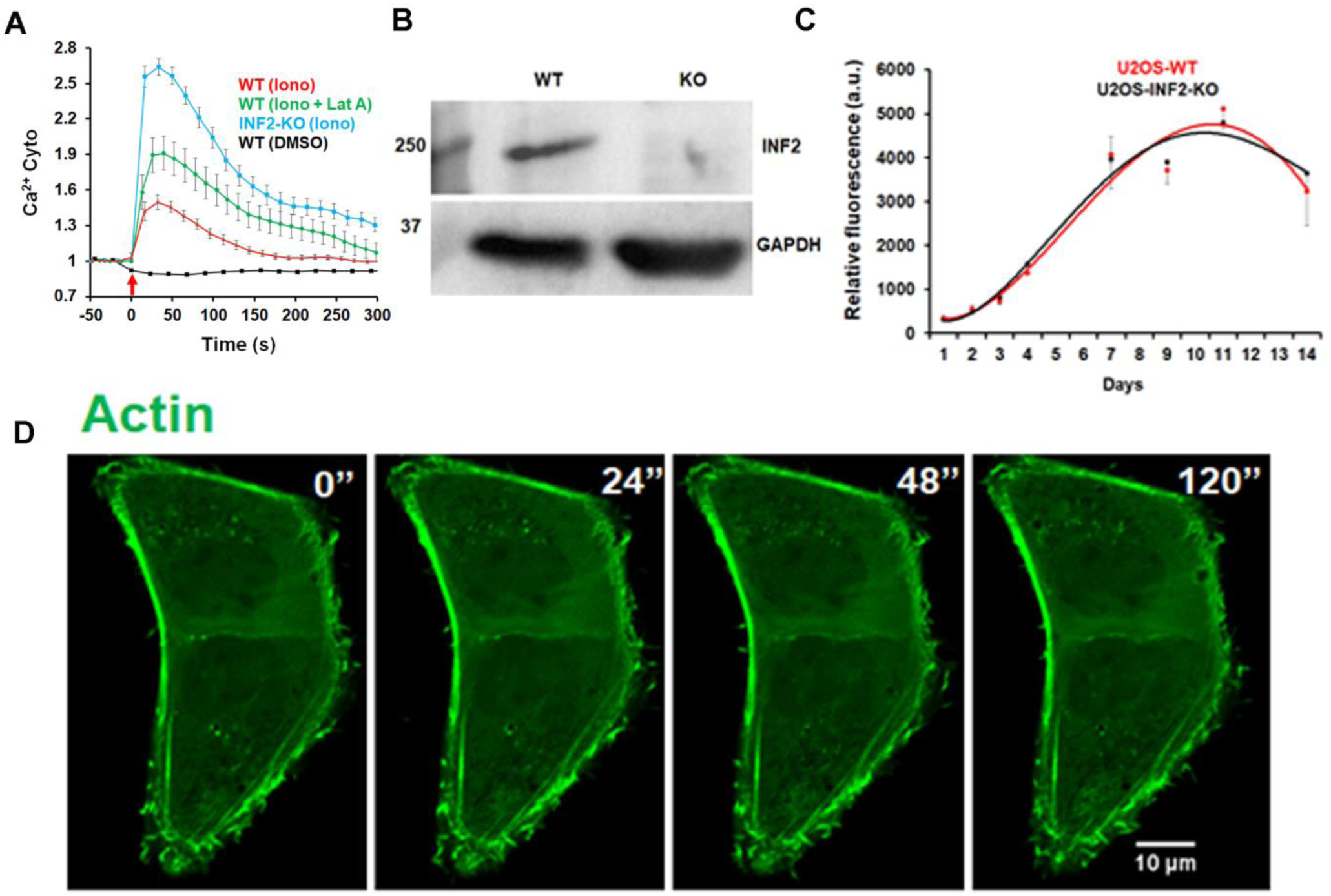
LatA effects on cytosolic calcium, and INF2-KO U2OS cell characterization. A) Quantification of cytoplasmic calcium spike following ionomycin (4 μM) treatment in U2OS-WT or INF2-KO cells, either with or simultaneous LatA treatment (2 μM). Ionomycin treatment at time 0 (arrow). N=10-15 cells. Error bars, SEM. B) Western blot analysis of U2OS-WT and U2OS-INF2-KO cell lines probed with anti-INF2 (upper) and anti-GAPDH (lower). C) Cell proliferation assay (Alamar blue) of U2OS-WT or U2OS-INF2_KO cells. Three replicates taken for each time point (median shown, with error bars representing minimum and maximum). Starting density: 5000 cells/well (24-well plate). Representative result from two independent experiments. D) Representative confocal image montage of U2OS-INF2-KO cells transfected with GFP-Ftractin (filamentous actin, green) and stimulated with 4 μM ionomycin (in the presence of 10 % serum) at time 0. Time in sec.

**Figure S4.**
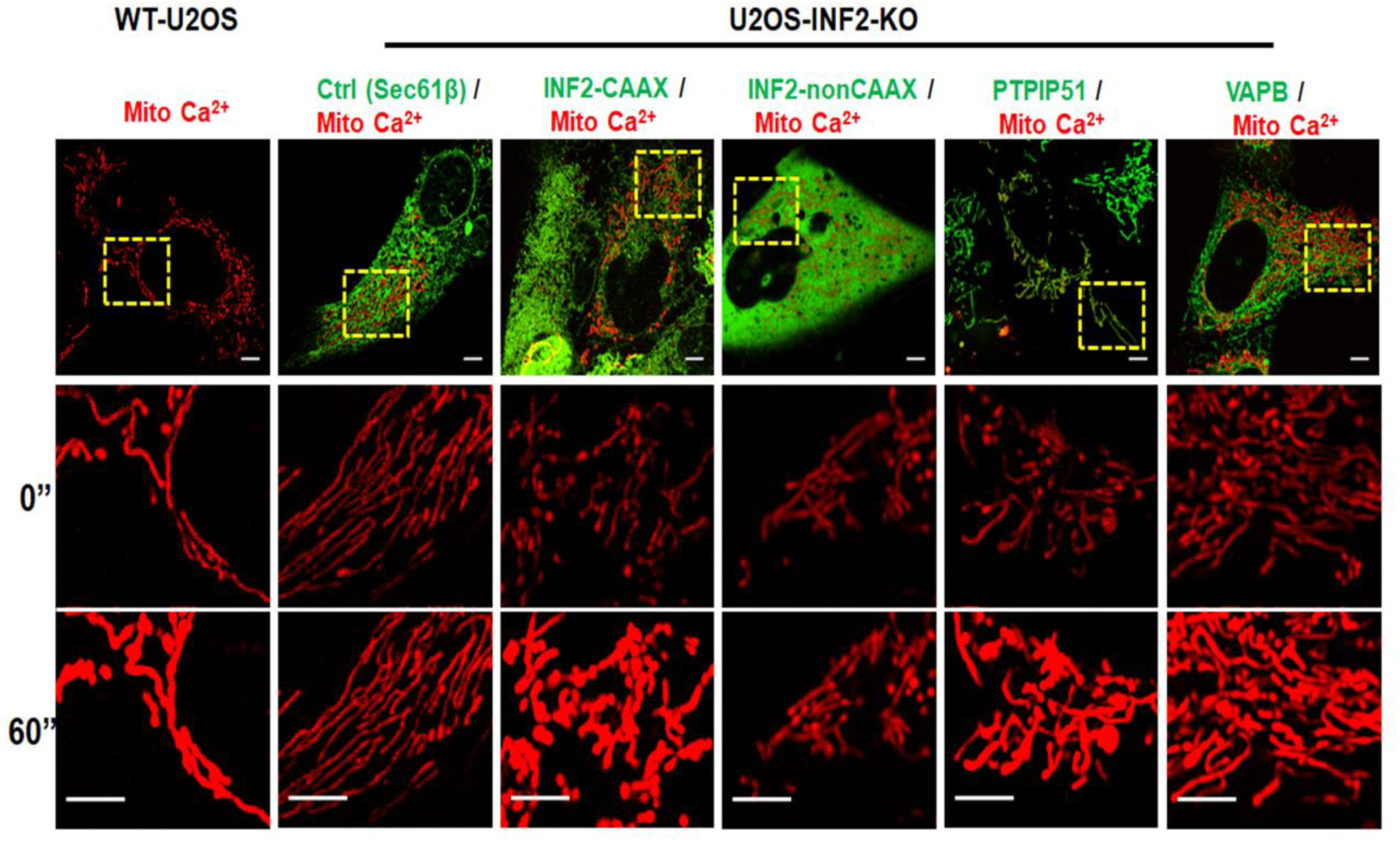
Rescue of mitochondrial calcium response in INF2-KO cells by over-expression of ER-mitochondrial tethers. INF2-KO U2OS cells were transfected with a mitochondrial matrix calcium probe (Mito-R-GECO) along with either CFP-VAPB or GFP-PTPIP51, then stimulated with 4 μM ionomycin. The effects of either GFP-INF2-CAAX or GFP-INF2-nonCAAX re-expression are shown for comparison. Insets at 0” and 30” after ionomycin stimulation. Scale bar: 5 μm (main panel); 2 μm (Insets).

**Figure S5.**
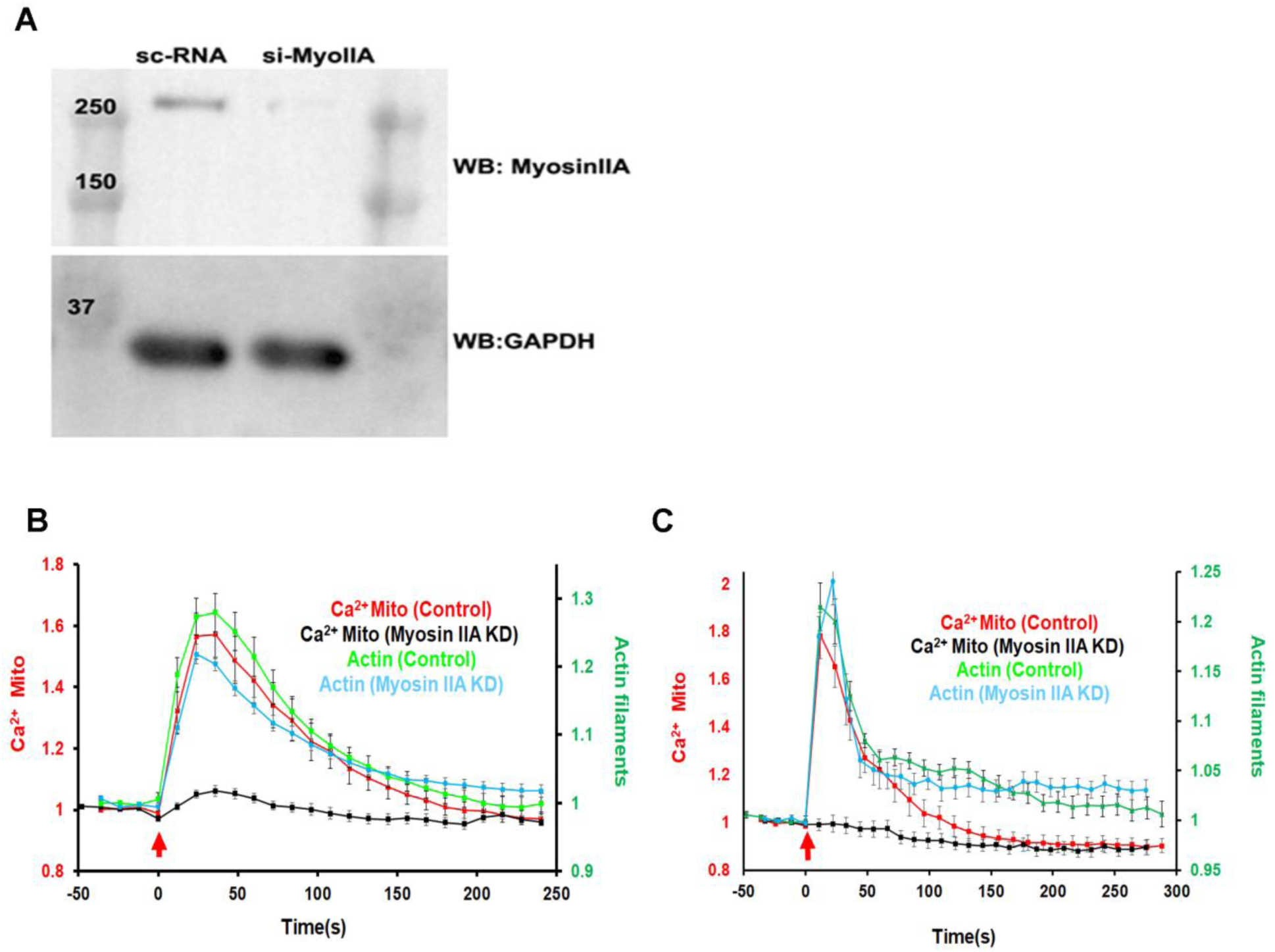
Effect of myosin IIA suppression on mitochondrial calcium spike. U2OS cells were transfected with siRNA against myosin IIA for 72 hrs, then transfected with mito-R-GECO and GFP-Ftractin. **A:** Western blot analysis of U2OS-WT and U2OS-Myosin IIA KD cells probed with anti-Myosin IIA (upper) and anti-GAPDH (lower). **B:** Actin polymerization burst and mitochondrial calcium spike following 4 μM ionomycin treatment **C:** Actin polymerization burst and mitochondrial calcium spike following 100 μM histamine treatment.

**Figure S6.**
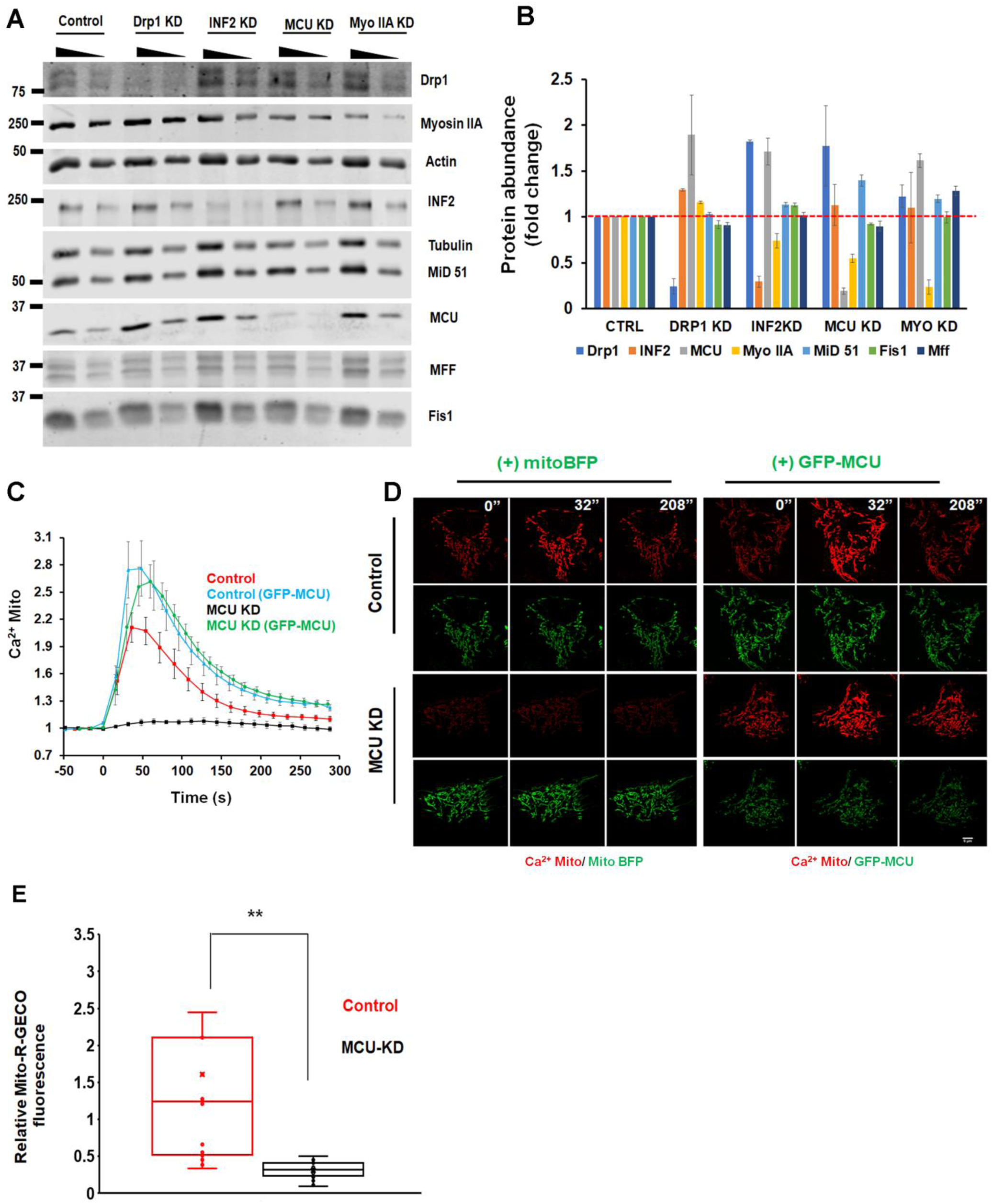
**A)** Western blot analysis of protein levels for U2OS cells transfected with siRNAs against MCU, Drp1, INF2, and myosin IIA. Proteins analyzed also included the mitochondrial division factors Mff, MiD51 and Fis1. Wedges refer to the two volumes of extract loaded on the gel (100% and 50%). **B)** Quantification of protein abundance from western blot analysis. Data from two independent experiments consisting of two volumes of loaded cell extract analyzed. Abundance in each knock-down was normalized to the control cell value. **C)** Rescue of ionomycin-induced mitochondrial calcium spike by expression of GFP-MCU. U2OS cells were transfected with either scrambled siRNA (control) or MCU siRNA (MCU KD) for 72 hrs, then transfected with mito-R-GECO with either mito-BFP or GFP-MCU. After 24 hrs, cells were stimulated with 4 μM ionomycin and the mitochondrial calcium spike was measured. **D)** Timelapse montages of experiment quantified in panel B. Green represents mito-BFP (left) or GFP-MCU (right). Red represents mito-R-GECO. Scale bar: 5μm **E)** Quantification of basal mito-R-GECO signal for un-stimulated control or MCU KD cells. Signal of mito-R-GECO was normalized to signal for co-transfected mito-BFP. N= 16 cells (Control); 17 cells (MCU KD). Each point represents one cell.

**Figure S7.**
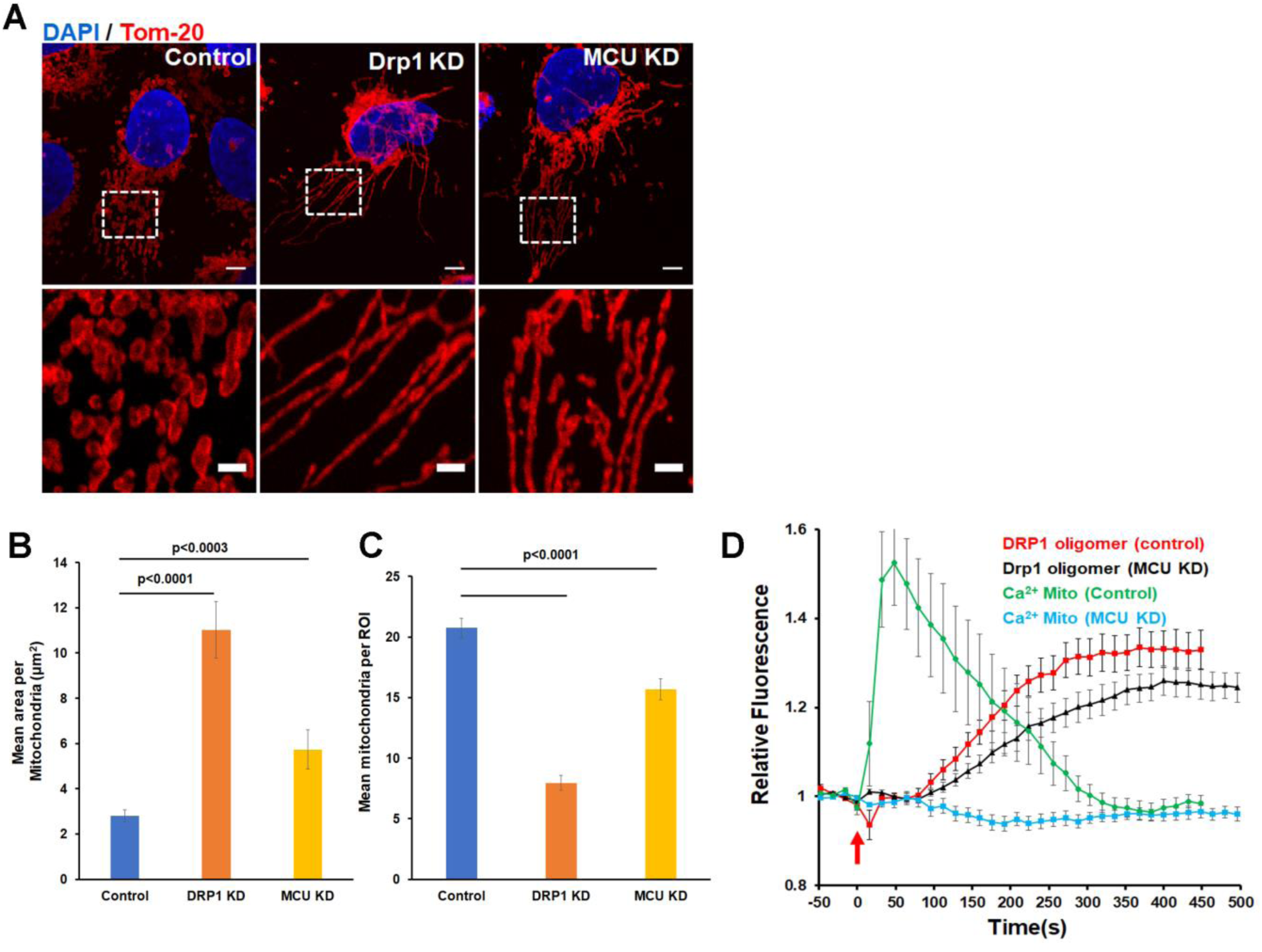
Assessment of mitochondrial division deficiency in MCU KD cells by fixed-cell analysis of mitochondrial size. U2OS cells were transfected with scrambled siRNA (Control), MCU siRNA (MCU KD), and Drp1 siRNA (Drp1 KD) for 72 hrs. Cells were then fixed and mitochondria stained using anti-Tom20 (red) and DAPI (blue). ROIs of fixed dimension were analyzed for mitochondrial length and number as described in the Materials and Methods. A) Images of Control (left), Drp1 KD (middle), or MCU KD cells (right). B) Mean mitochondrial length quantification represented as area (μm^2^) per mitochondrion for 80, 53, and 50 cells for Control, MCU KD, and Drp1 KD cells, respectively. Errors: SEM C) Mean mitochondrial number quantification, from same data set as in B. Errors: SEM. D) Drp1 oligomerization kinetics in MCU KD cells, as measured in whole cell. GFP-Drp1-knockin U2OS cells were treated with control siRNA or MCU siRNA for 72 hrs, then stimulated with 4 μM ionomycin while acquiring GFP images. The amount of oligomerized Drp1 was assessed as GFP signal above the cytoplasmic background signal, as described in Methods. Error bars: SEM.

**Figure S8.**
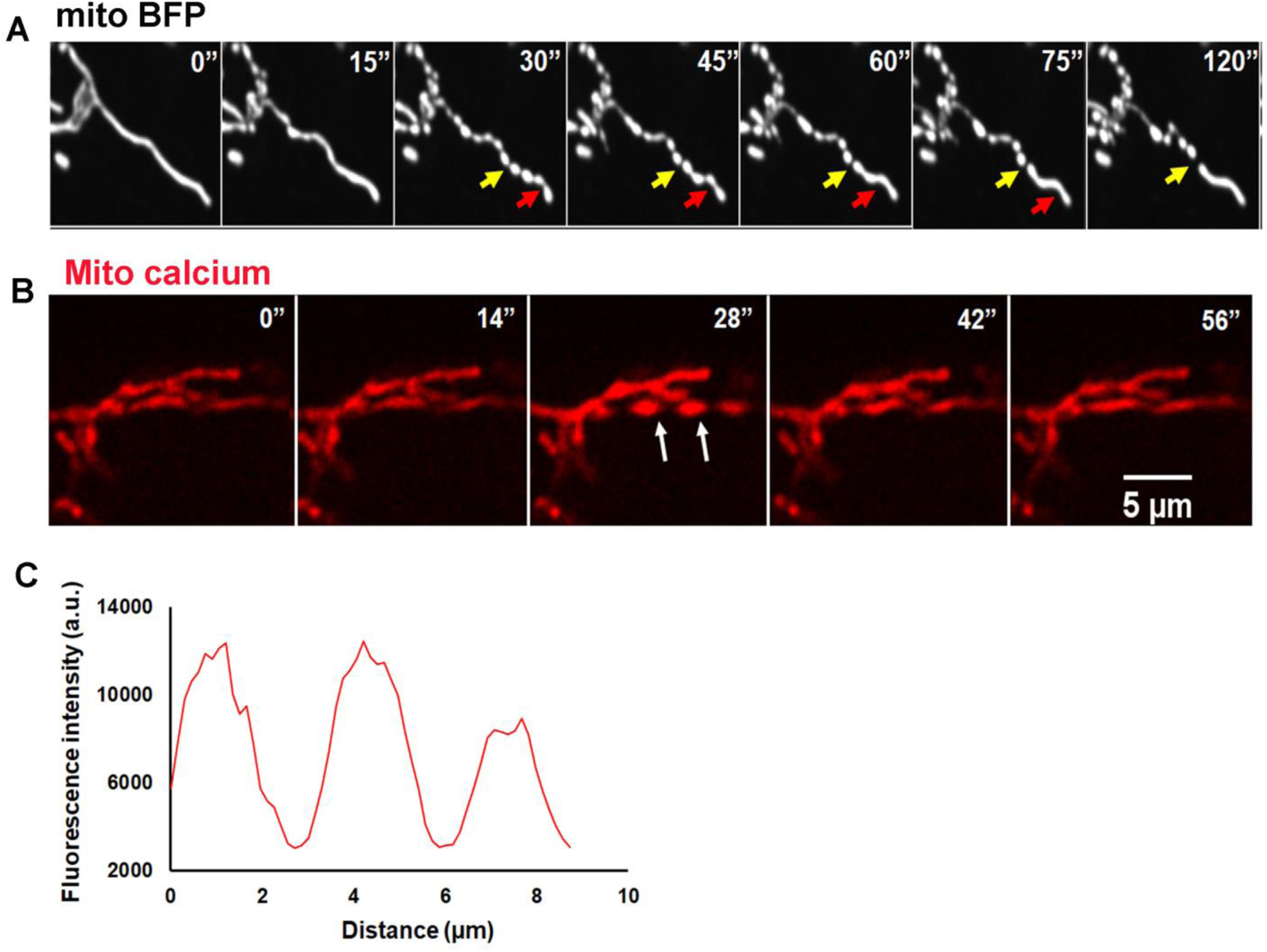
Spontaneous mitochondrial constrictions corresponding to increased mitochondrial calcium. **A**: Confocal image montage of spontaneous mitochondrial constrictions in U2OS cell transfected with mitoDsRed construct. Arrows: red - constriction that relaxes back without dividing; yellow - constriction resulting in division. Time in sec. **B**: Representative confocal image montage of spontaneous mitochondrial constrictions in U2OS cell transfected with mito-R-GECO (mitochondrial calcium) construct showing calcium spike associated with constriction (arrow). Time in sec. **C**: Linescan showing periodicity of spontaneous constrictions from U2OS cells transfected with mito-R-GECO construct.

**Figure S9.**
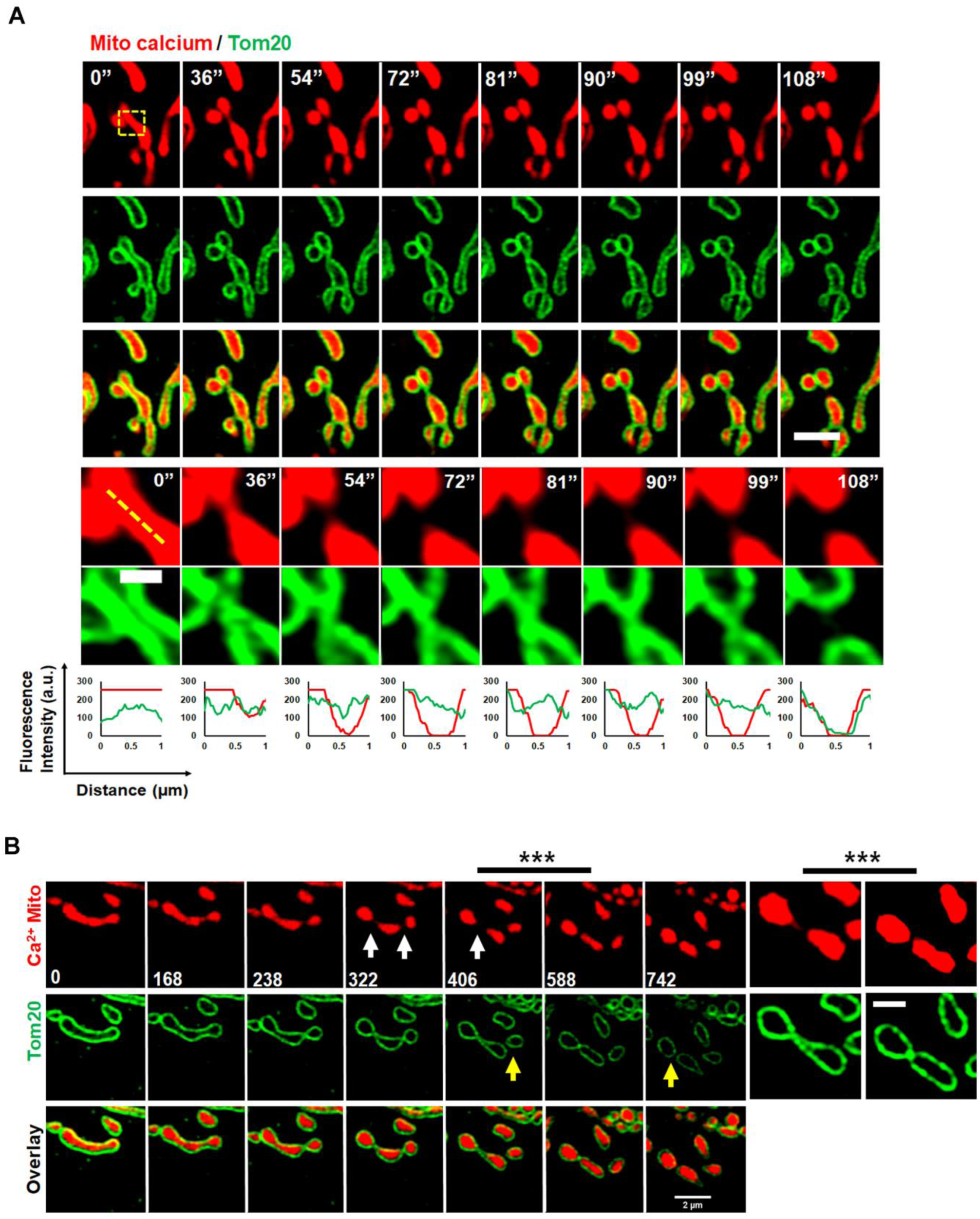
Matrix division prior to OMM division in U2OS cells. **A:** Second example of phenomenon shown in Figure 8A: matrix division prior to OMM division after ionomycin stimulation of U2OS cells transfected with mito-R-GECO (mitochondrial matrix calcium, red) and GFP-Tom20 (OMM, green). Scale Bar: Main panel: 2 μm; Inset: 0.5 μm **B:** Example of matrix division prior to OMM division after CGP37157 treatment of U2OS cells transfected with mito-R-GECO (mitochondrial matrix calcium, red) and GFP-Tom20 (OMM, green). White arrows show matrix division events, and yellow arrow show the subsequent OMM division event. Images at the right show two of these time points (***) as zoomed and enhanced images for mito-R-GECO, in an attempt to detect a lingering matrix tether. Scale bars: Main panels: 2 μm; Zoomed panels: 1 μm. **Video 10.**

**Figure S10.**
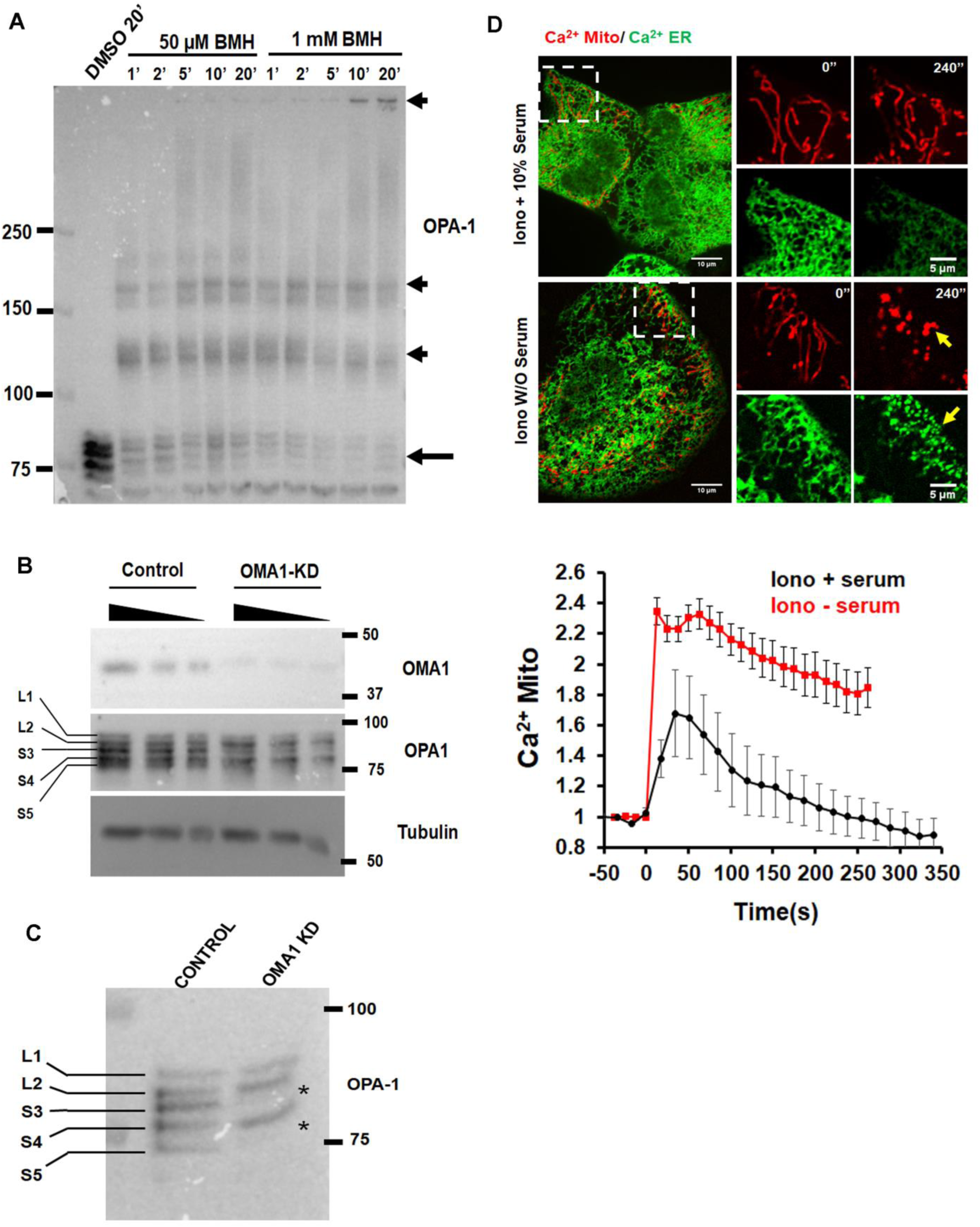
Opa1 and Oma1 experiments. **A)** BMH crosslinking assay to examine Opa1 oligomers. Control experiment to test time and concentration of BMH. Crosslinker was added to live U2OS cells in medium for the indicated times, then cells were lysed by medium removal and addition of SDS-containing buffer. Samples were analyzed by anti-Opa1 western blot. **B)** Western blot of U2OS transfected with siRNA for Oma1. Top panel shows Oma1 depletion. Middle panel shows depletion of Opa1 S3 and S5 bands, and increase in L2 and S4 bands, upon Oma1 depletion, as shown in Anand et al, 2014. **C)** Western blot of Opa1 bands in control and Oma1 KD cells at high resolution, showing depletion of S3 and S5 bands, and increases in L2 and S4 bands upon Oma1 depletion. **D)** Comparison of ionomycin stimulation in the presence or absence of serum. Top: representative confocal images of U2OS cells transfected with mito-R-GECO (mitochondrial calcium, red) and ER-GCaMP6-150 (endoplasmic reticulum calcium, green), stimulated with 4 ionomycin in the absence or presence (10%) of serum at time 0. Time in sec. Yellow arrows indicate fragmented mitochondria and ER. Bottom: quantification of mitochondrial calcium spike following 4 μM ionomycin treatment in the absence or presence of 10% serum in U2OS cells transfected with mito-R-GECO and treated at time 0. N=10 cells. Error bars, SEM.

**Video 1.** Ionomycin-stimulated increases in cytoplasmic calcium (Cyto-R-GECO), actin polymerization (GFP-Ftractin), and mitochondrial calcium (Mito-R-GECO) in U2OS cells (4 μM ionomycin). Movie at 5 frames/sec showing the first 25 seconds. Arrows indicate the incidence of each of the events. Corresponds to graphs in Figure 1B.

**Video 2.** Histamine-stimulated increases in cytoplasmic calcium (Cyto-R-GECO), actin polymerization (GFP-Ftractin), and mitochondrial calcium (Mito-R-GECO) in U2OS cells (100 μM histamine). Movie at 5 frames/sec showing the first 15 seconds. Arrows indicate the incidence of each of the events. Corresponds to graphs in Figure 1E.

**Video 3.** Ionomycin-stimulated actin burst (GFP-Ftractin) and mitochondrial calcium spike (Mito-R-GECO) in the same cell. Movie at 1 frame/sec. Corresponds to Figure 1G.

**Video 4.** Histamine-stimulated actin burst (GFP-Ftractin) and mitochondrial calcium spike (Mito-R-GECO) in the same cell. Movie at 2 frame/sec. Corresponds to Figure 1H.

**Video 5.** Effects of DMSO, ionomycin (4 μM), or histamine (100 μM) on ER calcium release. U2OS cells transfected with (ER-GCamP6-150) and treated with either of the above-mentioned drugs as indicated. Corresponds to Figure 2A.

**Video 6.** Ionomycin-induced actin morphology in INF2-KO cells re-expressing GFP-INF2-CAAX or GFP-INF2-nonCAAX. Cells transfected with INF2 construct, actin marker (GFP-Ftractin) and ER marker (ER-RFP) and treated with 4 μM ionomycin. Arrows indicate actin filaments on ER (INF2-CAAX) and non-ER regions (INF2-nonCAAX). Blue arrowhead denotes actin filament formation on the nuclear envelope for INF2-CAAX. Corresponds to Figure 3F.

**Video 7.** Comparison of ionomycin-induced mitochondrial matrix constrictions in WT, Drp1-KD or MCU-KD U2OS cells, using Mito-R-GECO (mitochondrial matrix calcium-sensor, red, left) and MitoBFP (matrix marker, blue, right). Corresponds to Figure 7B.

**Video 8.** Mitochondrial constrictions at ER contact sites. U2OS cell transfected with GFP-Sec61β (green, ER marker) and mito-R-GECO (red, mitochondrial calcium) and imaged live during ionomycin stimulation (4 μM). Corresponds to Figure 7C.

**Video 9.** Matrix division prior to OMM division during ionomycin-stimulated mitochondrial division. U2OS cells transfected with mito-R-GECO (mitochondrial matrix calcium, red) and GFP-Tom20 (OMM, green). Cells imaged live after addition of ionomycin (4 μM). Arrow indicates apparent matrix separation and arrowhead indicates OMM separation. Corresponds to Figure 8A.

**Video 10.** Matrix division prior to OMM division during CGP37157 (80 μM)-stimulated mitochondrial division. U2OS cells transfected with mito-R-GECO (mitochondrial matrix calcium, red) and GFP-Tom20 (OMM, green). Arrows indicate apparent matrix separation and arrowheads indicate OMM separation. Corresponds to Figure S9B.

